# Transplantation of Human IPSC-derived Microglia Ameliorates Neuropathology and Circuit Dysfunction in Progranulin-Deficient Mice

**DOI:** 10.64898/2026.01.13.699312

**Authors:** Hayk Davtyan, Sarah Naguib, Yuliya Voskobiynyk, Jean Paul Chadarevian, Joia K. Capocchi, Jeannie L. Giacchino, Ghazaleh Eskandari-Sedighi, Vivianna DeNittis, Jeremy B. Ford, Ani Agababian, Jasmine Nguyen, Alina L. Chadarevian, Madison S. Sutherland, Sepideh Kiani Shabestari, Alissa L. Nana, Jiasheng Zhang, Salvatore Spina, Man Ying Wong, Lea T. Grinberg, William W. Seeley, Eric Huang, Claire D. Clelland, Shiaoching Gong, Li Fan, Jeanne T. Paz, Mathew Blurton-Jones, Li Gan

## Abstract

Frontotemporal dementia (FTD) is a major cause of early-onset neurodegeneration characterized by progressive behavioral, emotional, and cognitive decline. Progranulin haploinsufficiency, a leading genetic cause of familial FTD, disrupts lysosomal function, lipid metabolism, autophagy, and neuroimmune signaling across multiple cell types. Increasing evidence indicates that microglia are particularly sensitive to progranulin loss, exhibiting elevated complement activation that contributes to TDP-43 proteinopathy and neuronal dysfunction. Here, we investigate the biological role of restoring progranulin exclusively within microglia by transplanting human induced pluripotent stem cell-derived microglia (iMG) into progranulin (*Grn*)-deficient mice. We find that wild-type, but not *Grn*-deficient, human iMG restore brain-wide progranulin levels, normalize microglial transcriptional states, and ameliorate pathological, functional, and behavioral phenotypes associated with progranulin loss. Because microglia are the only source of progranulin in this system, these findings demonstrate that microglial progranulin is sufficient to restore key aspects of cellular, circuit, and behavioral homeostasis in a progranulin-deficient FTD model. More broadly, this work highlights a central, microglia-intrinsic role for progranulin in maintaining brain function and provides a framework for dissecting microglia-specific mechanisms across FTD and related neurodegenerative disorders.

**One Sentence Summary:** Our study demonstrates that xenotransplantation of wild-type human iPSC-derived microglia into progranulin-deficient mice mitigates core neuropathological, network-level, and behavioral features of Frontotemporal Dementia.

## INTRODUCTION

Frontotemporal dementia (FTD) is the second most common cause of dementia in individuals under 65, and is characterized by the progressive development of behavioral, emotional, and cognitive impairments^1–3^. Despite its prevalence, there are currently no effective treatments for FTD, highlighting a critical need to understand the mechanisms that underlie this disease to develop effective therapeutic approaches. Among the known genetic contributors to FTD, progranulin (human gene symbol: *GRN*) haploinsufficiency is the second most prevalent cause of familial FTD^4–6^.

In the brain, *GRN* is most highly expressed by microglia, yet progranulin deficiency disrupts essential cellular functions across multiple cell types, including disruptions in lysosomal activity, lipid metabolism, autophagy, and neuroimmune signaling ^4,5,7–9^. Recent evidence highlights that microglial dysfunction is a key contributor to progranulin-deficient FTD; microglia in FTD display reduced phagocytic efficiency, impaired motility, and elevated production of complement, which is associated with neuronal TDP-43 proteinopathy and neurodegeneration^10–14^. Moreover, microglia-specific deletion of *Grn* is sufficient to induce neuronal hyperexcitability in a *Grn*-deficient mouse model^15,16^; notably, the excessive grooming phenotype characteristic of *Grn*-deficient mice is prevented by inactivation of NF-κB specifically in microglia^13^. Together, these findings underscore that progranulin plays a critical, microglia-intrinsic role in maintaining cellular homeostasis and regulating neuroimmune function. Rather than simply serving as a global neuroprotective factor, progranulin appears essential for preserving key microglial processes, including lysosomal function, lipid handling, and inflammatory signaling, all of which are disrupted in FTD and are increasingly implicated across other neurodegenerative diseases. Understanding how progranulin deficiency impacts microglial biology and its influence on neuronal vulnerability is, therefore, fundamental for elucidating FTD mechanisms, and may reveal broader principles of neuroimmune dysfunction relevant to Alzheimer’s disease, Parkinson’s disease, and related disorders.

To determine whether microglial loss of progranulin is a causal driver of frontotemporal dementia (FTD) pathogenesis in a human-relevant context, we transplanted wildtype human iPSC-derived microglia into xenotolerant *Grn^−/−^* mice, thereby achieving microglia-restricted restoration of progranulin in vivo. *Grn^−/−^* mice exhibit lysosomal dysfunction, lipofuscin accumulation, microgliosis, and mild behavioral deficits that closely recapitulate the neuropathological and behavioral features reported in Grn-null models^17–19^. For simplicity, *Grn^−/−^*mice are hereafter referred to as *Grn^−/−^* throughout the manuscript. Rather than broadly elevating progranulin levels throughout the brain, this approach allowed us to directly assess whether correction in this single cell type is sufficient to ameliorate disease. Using integrated biochemical, histological, and transcriptional analyses together with slice electrophysiology and behavioral assays, we show that engraftment of wild-type, but not progranulin-deficient, iMG, robustly attenuates the neuropathological, circuit-level, and behavioral abnormalities associated with progranulin deficiency. These results establish microglia as a critical cellular locus of progranulin action: progranulin-deficient microglia fail to support neuronal health, whereas microglia with intact progranulin preserve key homeostatic and neuroimmune functions. Together, our findings demonstrate that restoring progranulin specifically in microglia is sufficient to confer broad, multi-level rescue, highlighting a central role for microglial progranulin in FTD.

## RESULTS

### Human microglia transplantation decreases lipofuscinosis in *Grn*-deficient mice

To directly test the role of human microglia-derived progranulin in vivo, we generated a progranulin-deficient xenograft model that permits stable engraftment of healthy human iPSC-derived microglia. By restricting progranulin restoration to microglia, this system isolates cell-autonomous microglial progranulin function while enabling functional assessment in a human-relevant in vivo context. *Grn^R^*^493^*^X/R^*^493^*^X^* mice, which model the most common human *GRN* mutation introducing a premature stop codon at arginine 493 (R493X), show markedly reduced *Grn* mRNA levels and complete loss of detectable progranulin protein^19^ and are referred to as *Grn^−/−^* throughout this study. These mice were backcrossed onto a *hCSF1/Rag2^−/−^/il2rg^−/y^*(hCSF1) background to generate xenotolerant *hCSF1-Grn^−/−^* recipients (**Figure 1A**). Immunoblotting confirmed near-complete loss of murine progranulin within the brain, liver, spleen, and plasma of the resulting *hCSF1-Grn^−/−^* mice (**Figure S1A**).

**Figure 1.**
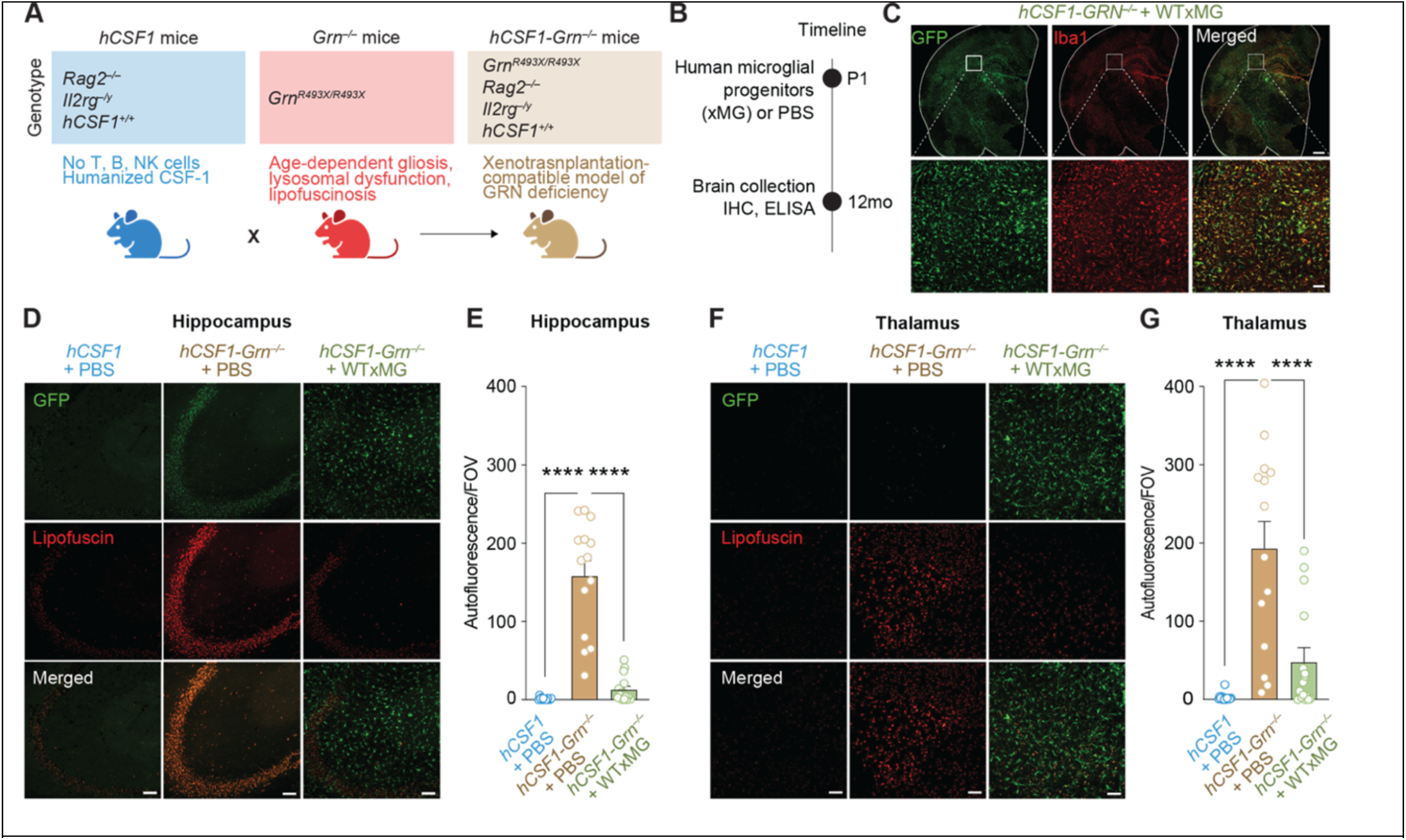
Human microglia transplantation decreases lipofuscinosis in Grn-deficient mice. **(A)** Xenotolerant *Grn*-deficient mice were generated by crossing immune-intact *Grn^−/−^*mice onto the humanized, immune-deficient *hCSF1^+/+^/Rag2^−/−^/Il2rg^−/y^*(*hCSF1*) knockin background and restoring homozygosity for all four alleles. **(B)** Schematic illustrating the neonatal transplantation and analysis paradigm for *hCSF1* and *hCSF1-Grn^−/−^* mice. **(C)** Representative confocal microscopy of xenografted GFP-positive (green) human microglia (Iba1, red) in 12-month-old *hCSF1-Grn^−/−^*+xMG adult mice; scale bars: 500 µm (low magnification), 50 µm (high magnification). **(D, F**) Representative confocal microscopy of human xMG (GFP, green) and autofluorescent inclusions (lipofuscin, red) in the hippocampus **(D)** and thalamus **(F)** of WT and transplanted mice; scale bar, 50 µm. **(E, G)** Quantification of autofluorescence within the hippocampus **(E)** and thalamus **(G)** of wild-type and transplanted mice; P values from one-way ANOVA (Hippocampus: F_2,42_ = 65.00; ^∗∗∗∗^*p* < 0.0001; Thalamus: F_2,42_ = 20.86; ^∗∗∗∗^*p* < 0.0001) with Tukey’s multiple comparisons tests. Data represented as mean value ± SEM (non-injected *hCSF1* littermates, n = 15; *hCSF1-Grn^−/−^* +PBS, n = 14; *hCSF1-Grn^−/−^* +xMG, n = 16) (Female, n = 8 per group; Male, n = 6–8 per group). ns, not significant; ^∗∗∗∗^*p* < 0.0001.

Next, neonatal *hCSF1-Grn^−/−^* mice were injected with either phosphate buffered saline control (*hCSF1-Grn^−/−-^* +PBS) or xenografted with wild-type (WT) GFP-expressing human microglial progenitors (*hCSF1-Grn^−/−-^* +WTxMG). Wild-type hCSF1 littermates were similarly injected with PBS (*hCSF1* +PBS). Mice were then aged to 12 months for all subsequent analyses, as previous studies have indicated that progranulin deficiency causes an age-dependent upregulation in lysosomal and proinflammatory genes in microglia^8^ (**Figure 1B**). Immunohistochemistry (IHC) for ionized calcium-binding adapter molecule 1 (Iba-1), a commonly used microglial marker^22^, and green fluorescent protein (GFP) revealed widespread engraftment of human xenografted microglia (xMG) in 12-month-old *hCSF1-Grn^−/−^* +WTxMG mice (**Figure 1C**). To determine whether the transplantation of healthy human microglia can elevate levels of progranulin, soluble brain extracts were examined using ELISAs for human or mouse progranulin. As expected, levels of human progranulin were significantly elevated in *hCSF1-Grn^−/−^* +WTxMG engrafted mice, and analysis with a mouse-targeted ELISA further confirmed the loss of mouse progranulin in *hCSF1-Grn^−/−^* +PBS brains (**Figure S1E, F**). Owing to partial cross-reactivity with human progranulin, the murine-targeted ELISA detected a modest but significant increase in brain progranulin levels in hCSF1-*Grn^−/−^*+ WT xMG mice (**Figure S1F**). Thus, transplantation of wild-type human microglia elevates brain progranulin levels in *Grn*-deficient mice.

Previous studies have shown that *Grn^−/−^* mice develop lysosomal dysfunction and lipofuscinosis, a neuronal pathology commonly observed in FTD postmortem brains^17,20^. We therefore examined unstained sections from all three groups by confocal microscopy as previously described^21–23^ to quantify the accumulation of autofluorescent lipofuscin within the brains of *hCSF1* +PBS littermates and *hCSF1-Grn^−/−^* mice engrafted with PBS or xMG. This analysis revealed significantly elevated levels of autofluorescent inclusions indicative of lipofuscinosis within the hippocampus and thalamus of *hCSF1-Grn^−/−^* +PBS mice in comparison to *hCSF1* littermates (**Figure 1D–G**). In contrast, the presence of progranulin restricted to microglia significantly reduced lipofuscin levels in 12-month-old mice, restoring autofluorescence levels to those of *hCSF1* controls.

### Xenotransplanted human microglia reduce markers of inflammation and microglial activation in *hCSF1-Grn^−/−^* mice

In progranulin-deficient FTD patients, microglial activation in the frontal cortex predicted cognitive decline^11^. *Grn^R^*^493^*^X/R^*^493^*^X^* mice phenocopy the microgliosis and hyperinflammatory phenotype of *Grn*^−/−^ mice within the CNS^17,19,24–27^. To evaluate the impact of wildtype microglial transplantation on neuroinflammation, we measured the levels of proinflammatory cytokines and chemokines in soluble brain extracts from both WT littermates and transplanted *hCSF1-Grn^−/−^* mice. A multiplex ELISA assay revealed that *hCSF1-Grn^−/−^*+PBS mice exhibited significantly elevated levels of several proinflammatory cytokines, including IL-1β, TNF-α, IL-33, IP-10, and MIP-1α, within the brain. Importantly, engraftment of wild-type human microglia prevented their elevation, maintaining cytokine levels comparable to those of *hCSF1* controls (**Figure 2A**).

**Figure 2.**
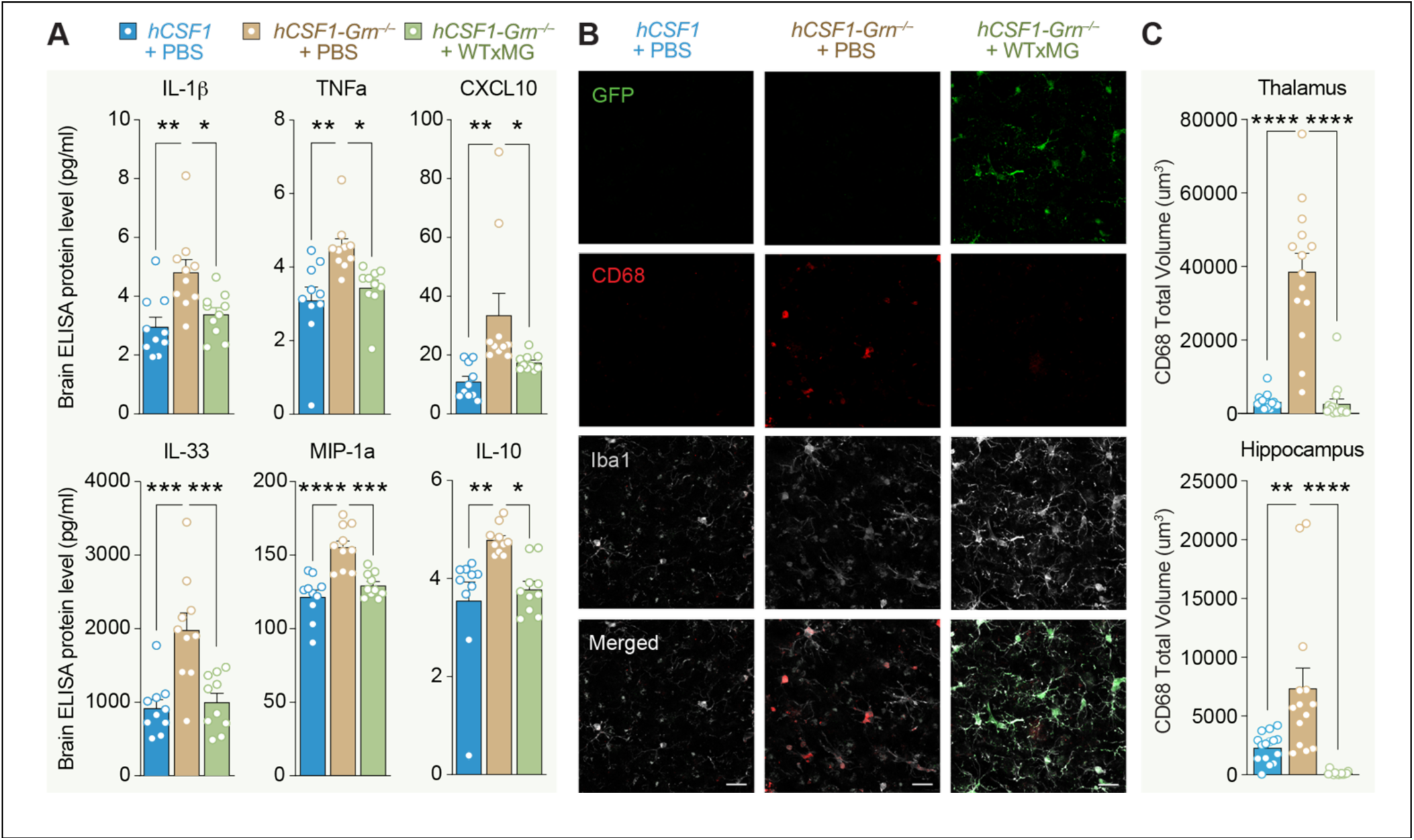
Human microglia engraftment reduces markers of inflammation and microglial activation in *hCSF1 - Grn^R^*^493^*^X/R^*^493^*^X^* mice. **(A)** Quantification of proinflammatory IL-1*β*, TNF-*⍺*, CXCL10, IL- 33, MIP-1a, and IL-10 levels in soluble brain extracts of *hCSF1* littermates and *hCSF1-Grn^−/−^* + PBS and *hCSF1-Grn^−/−^* + WTxMG mice; *p-*values from one-way ANOVA (IL-10: *F*_2,27_ =7.332; \*\**p* = 0.0029) (IL-1B: *F*_2,27_ = 7.958; \*\**p* = 0.0019) (TNF-a: *F*_2,27_ = 7.445; \*\**p* = 0.0027) (IL-33*: F*_2,27_ = 12.78; \*\*\**p* = 0.0001) (IP-10: *F*_2,27_ = 6.734; \*\**p* = 0.0042) (MIP-1a: *F*_2,27_ = 18.61; \*\*\*\**p* < 0.0001) with Tukey’s multiple comparisons tests. Data represented as average mean value ± SEM (*hCSF1* littermates, *n* = 10; *hCSF1-Grn^−/−^* +PBS, *n* = 10; *hCSF1-Grn^−/−^* +WTxMG, *n* = 10) (Female, *n* = 5 per group; Male, *n* = 5 per group). **(B)** Representative confocal microscopy of human xMG (GFP, green), microglial marker (IBA1, white), and microglial activation marker (CD68, red) in the thalamus (top) and hippocampus (bottom) of wild-type and transplanted *Grn^−/−^* mice, scale bar, 50 µm. **(C)** Quantification of CD68-positive immunoreactivity (total volume/FOV) within the thalamus (top) and hippocampus (bottom) of wild-type and transplanted mice *p* values from one-way ANOVA (Hippocampus: *F*_2,42_ = 24.32\*\*\*\**p* < 0.0001; Thalamus: *F*_2,42_ = 17.43; \*\*\*\**p* < 0.0001) data analyzed with Tukey’s multiple comparisons tests. Data are presented as mean ± SEM (*hCSF1* littermates, *n* = 15;*hCSF1-Grn^−/−^* +PBS, *n* = 14; *hCSF1-Grn^−/−^* + WTxMG, *n* = 16) (Female, *n* = 8 per group; Male, *n* = 6–8 per group). ns, not significant, ^∗^*p* < 0.05, ^∗∗^*p* < 0.01, ^∗∗∗^*p* < 0.001, and ^∗∗∗∗^*p* < 0.0001. Comparisons not shown are not significant.

Surprisingly, IL-10, an anti-inflammatory cytokine, was similarly elevated in *hCSF1-Grn^−/−^* +PBS mice but reduced with microglial engraftment, suggesting that changes in IL-10 may represent a compensatory attempt to dampen the chronically heightened proinflammatory state of *Grn^−/−^* brains (**Figure 2A**). The effects of xenografts were specific to these six cytokines as levels of IL-9, IL-15, IL-17A/F, IL-27p/28 (IL-30), and MIP-2 (CXCL2) remained elevated in *hCSF1-Grn^−/−^* +xMG brains (**Figure S2A–D**). Five other analytes (IL-12p70, IL-2, IL-6, KC/GRO, and MCP-1) showed no significant differences between groups (**Figure S2E–I**).

Next, we analyzed immunoreactivity of CD68, a microglial phagolysosome membrane protein commonly used as a marker of microglial activation^25,26^ and previously established as a binding partner of progranulin and progranulin-derived granulin E^28^. In the thalamus, the observed increase in CD68 immunoreactivity within *hCSF1-Grn^−/−^* + PBS mice was prevented, and CD68 was maintained at WT levels in *hCSF1-Grn^−/−^* +xMG mice (**Figure 2B, C)**. Similarly, CD68 immunoreactivity in the hippocampus was significantly lower in *hCSF1-Grn^−/−^* +xMG mice in comparison to both *hCSF1* controls or *hCSF1-Grn^−/−^*

+PBS mice (**Figure 2B, C**). Together, these results demonstrate that progranulin derived from human wild-type microglia is sufficient to reduce neuroinflammation and microgliosis in progranulin-deficient mice.

### Transcriptional signatures of *GRN^−/−^* vs. *GRN^+/^*^+^ xMG in *hCSF1-Grn^−/−^* mice

To further dissect the protective effects of microglial progranulin, we xenotransplanted *hCSF1-Grn^−/−^* neonatal mice with isogenic human *GRN^+/+^* (WTxMG) or *GRN^−/−^* (KOxMG) microglial progenitor cells (**Figure 3A**). At 9.5 months, we performed behavioral analyses to assess grooming behaviors. Mice were then sacrificed *hCSF1-Grn^−/−^* at 10.5–11.5 months to conduct electrophysiology and collect hippocampal and thalamic tissue for single-nuclei RNA sequencing (snRNA-seq) (**Figure 3B**). We also performed biochemical analysis of soluble extracts prepared from the remaining brain tissue to confirm the absence of progranulin protein in mice engrafted with KOxMG and robust progranulin expression in mice engrafted with WTxMG (**Figure 3C**).

**Figure 3.**
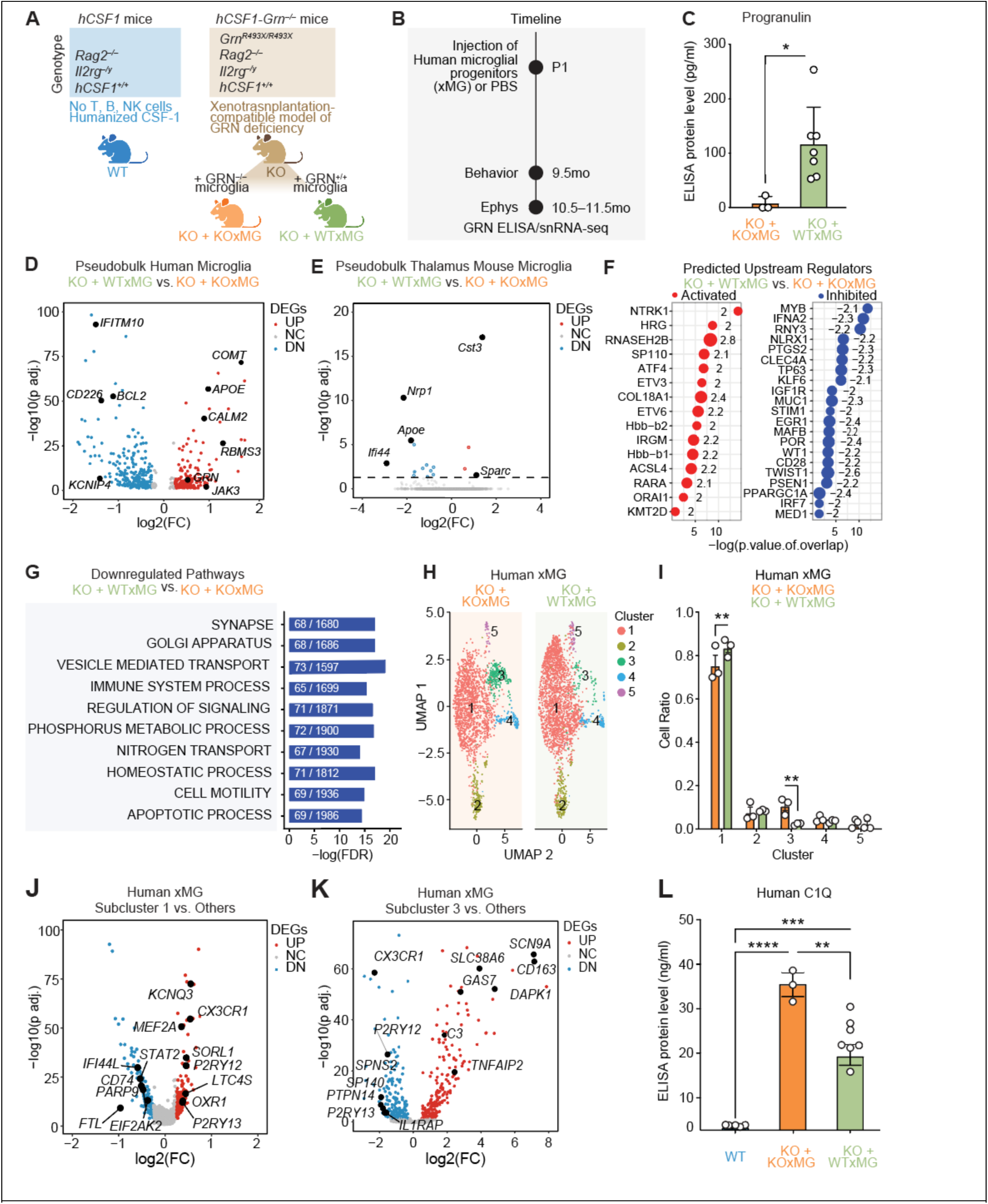
Transcriptional signatures of xenografted human KO-xMG and WT-xMG in hCSF1-Grn*^−/−^*brains. **(A)** Schematic of mouse strains: *hCSF1* littermates (WT) and *hCSF1-Grn^−/−^*xenografted with WT (WTxMG) or KO (KOxMG) human microglia. **(B)** Schematic experimental timeline of microglia injected at P1, behavior at 9.5 months, electrophysiology, and snRNA-seq at 10.5–11.5 months. **(C)** Quantification of human progranulin ELISA showing increased progranulin levels in WTxMG mice. Unpaired t-test (t(8) = 2.647, \**p* = 0.0294). Data represented as mean ± SEM (*hCSF1-Grn^−/−^* +KOxMG, n = 3; *hCSF1-Grn^−/−^* +WTxMG, n = 7). **(D)** Volcano plot of pseudobulk DEGs in human microglia between WTxMG and KOxMG groups, with key genes up- or downregulated. **(E)** Volcano plot of pseudobulk DEGs in thalamic mouse microglia between WTxMG and KOxMG groups, with key genes up- or downregulated. **(F)** Top predicted upstream regulators identified using Ingenuity Pathway Analysis (IPA). **(G)** Top predicted downregulated pathways in WTxMG v. KOxMG. **(H)** UMAP of human microglia by genotype, showing clustering differences between WTxMG and KOxMG groups. **(I)** Quantification of average cell ratios in each subcluster by group in human microglia. **(J)** Volcano plot of subcluster 1 DEGs in human microglia between WTxMG and KOxMG groups, with key genes up- or downregulated. **(K)** Volcano plot of subcluster 3 DEGs in human microglia between WTxMG and KOxMG groups, with key genes up- or downregulated. (**L**) Quantification of human C1Q ELISA showing significantly decreased levels in WTxMG injected mice. *p*-value from one-way ANOVA (F_2,12_=2.525, \*\*\*\**p <*0.0001) with Tukey’s multiple comparisons test. Data represented as mean ± SEM (*hCSF1*, n = 5*; hCSF1-Grn-/- +* KOxMG, n = 3*; hCSF1-Grn^−/−^ +* WTxMG, n = 7). \**p* < 0.05 and *\*\*p* < 0.01.

The transcriptomic profiles of human microglia were analyzed by isolating human cells from mouse cells *in silico* (**Figure S3, Table S1**). Mouse microglia accounted for 1–2% of total cells in samples that received human microglia injections, compared with ∼7% of total cells in non-injected mice, indicating robust engraftment and long-term persistence of xenografted human microglia at 10.5–11.5 months post-transplantation. In comparison to KOxMG, WTxMG expressed higher levels of *APOE*, *COMT*^29^, an enzyme involved in the breakdown of dopamine and norepinephrine, *CALM2*^30^, the critical calcium sensor known as calmodulin, and *RBMS3*^31^, a protein associated with maintaining cell health, supporting tissue repair, and promoting neuronal development (**Figure 3D**, **Table S1**). In contrast, WTxMG exhibited lower levels of several interferon-responsive genes, including *CD226, IFITM10*, *MX1*, and *IFI44L*^29^, and the antiapoptotic gene *BCL2*^32^ (**Figure 3D, Table S1**). As expected, *GRN* was among the upregulated genes in WTxMG (**Figure 3D**).

To assess how restoring progranulin specifically in transplanted human microglia reshapes the surrounding brain environment, we profiled endogenous mouse microglia isolated from the thalamus of mice receiving no graft (WT), GRN-deficient iMG (KO–KO), or wild-type iMG (KO–WT) (**Figure 3E**). Remarkably, only 19 differentially expressed genes (DEGs) were detected between the KO–WT and KO–KO groups, indicating that the *Grn^−/−^* background exerts a dominant constraint on the endogenous mouse microglial transcriptome. This minimal transcriptional divergence suggests that the broader molecular changes observed across our datasets are driven primarily by the transplanted human microglia rather than secondary reprogramming of host microglia. Notably, among the limited DEGs identified, several disease-associated and inflammatory markers, including *Apoe*, *Ifi44*, and *Nrp1*^33^, were downregulated in KO–WT mouse microglia, consistent with a modest but detectable anti-inflammatory influence of progranulin-replete human microglia on the local microglial niche.

Whereas endogenous mouse microglia showed minimal transcriptional remodeling, analysis of the human xenografted microglia (xMG) revealed robust engagement of specific regulatory programs. Ingenuity Pathway Analysis (IPA) further identified predicted activators of these differentially expressed genes, including *NTRK1*^34,35^ (TRK-A), which is essential for neuronal development and cholinergic neuronal survival, and *ETV3/ETV6,* implicated in monocyte differentiation^36^ (**Figure 3F**). Predicted upstream inhibitors included *IFNA2, RNY3*^37^, involved in modulating cell proliferation and *IL13* regulation, as well as *STIM1*, which regulates calcium entry and purinergic activation in microglia^38,39^ (**Figure 3F**). Gene set enrichment analysis also revealed that the top downregulated pathways in WTxMG are those involved in apoptosis, immune system processes, vesicle-mediated transport, and the regulation of signaling. (**Figure 3G**).

To define how progranulin status reshapes the transcriptional states of human xenografted microglia, we compared human microglia from KO–KO and KO–WT brains. Among the major human microglial clusters, subcluster 1 was significantly enriched in WTxiMG, whereas subcluster 3 was relatively expanded in KOxiMG (**Figure 3H, I**). Subcluster 1 displayed a canonical homeostatic profile, with preservation of microglial identity genes such as *P2RY12* and *CX3CR1*, *SORL1*^40^, and *KCNQ3*^41^, while key interferon/inflammatory mediators, including *STAT1* and *IFI44L,* were selectively downregulated compared with other human microglia (**Figure 3J**). Thus, the state that is favored in WTxMG is not only homeostatic but also actively anti-inflammatory, characterized by reduced expression of IFN/innate immune effectors.

By contrast, genes defining subcluster 3 were strongly upregulated in KOxMG, including activation and inflammatory markers as well as complement pathway components such as *IL1RAP*, *TNFAIP2*^42^, *CD163*^43^, *DAPK1*^44^, *GAS7*^45^, and *C3*, consistent with a more reactive, disease-associated microglial phenotype^46^ (**Figure 3K**). To validate that WTxMG reduces complement-mediated pathways, we performed a C1Q ELISA and found that restoring wild-type levels of progranulin in microglia was sufficient to normalize C1Q levels in *hCSF1-Grn^−/−^* mice (**Figure 3L**). Collectively, these data show that progranulin sufficiency in human microglia promotes a stable, anti-inflammatory homeostatic state, whereas progranulin deficiency drives an opposing inflammatory activation state. Consistent with this, engraftment of wild-type microglia elevated brain progranulin levels, ameliorated compensatory increases in microglial lysosomal gene expression observed in *Grn*-deficient mouse and human microglia⁸_˒_¹³, and restored a predominantly homeostatic microglial landscape.

### Differential effects of WTxMG and KOxMG on the transcriptomes of mouse astrocytes and oligodendrocytes

To directly test how progranulin produced by human microglia shapes non-cell-autonomous glial responses in vivo, we compared the effects of xenografted WTxMG or KOxMG on transcriptional states of murine glial cells within the hippocampus and thalamus, regions that are strongly linked to functional deficits in *Grn^−/−^* mice^5,27,47^. In the hippocampus, WTxMG downregulated astroglial *Pde10a* and upregulated *Inpp4b*, a gene involved in astrocyte-microglia communication^48^ (**Figure S3A, Table S2**). IPA of differentially expressed genes in hippocampal astrocytes revealed a coordinated shift in upstream regulatory programs in the KO-WT group compared to KO–KO controls, characterized by concurrent activation of trophic and survival-associated pathways and inhibition of stress- and degeneration-linked signaling (**Figure S3B, Table S2**). Upstream regulator analysis predicted increased activity of neurotrophic and pro-survival signaling nodes, including BDNF–NTRK2 signaling, mTOR–RPS6KB1-associated translational pathways^49^, and cAMP/CREB-linked signaling via GNAS^50^. In parallel, IPA predicted inhibition of multiple stress-responsive and degenerative regulators in KO-WT astrocytes, including ERN1- and RNASEL-linked stress pathways^51,52^, senescence- and lysosomal-associated regulators such as GLB1 and SPG21^53,54^, and immune-associated signaling, including CD24. Additional inhibition of metabolic and post-transcriptional regulators such as METTL3^55^ further supports a reduction in cellular stress and dysregulated immune signaling. Together, these findings indicate that progranulin produced by human microglia drives a shift in hippocampal astrocytes in *Grn^−/−^* mice toward a supportive, homeostatic state with reduced stress and degenerative signaling. Consistent with this shift, engraftment of progranulin sufficient human microglia (WTxMG) actively suppressed complement gene expression in KO astrocytes, whereas progranulin-deficient microglia (KOxMG) failed to restrain complement signaling (**Figure S3C**).

In the thalamus, WTxMG downregulated astroglial interferon-related *Ifi27*, as well as *C4b*, *Gfap*, and *Cxcl14*, genes associated with inflammation, complement pathways, and gliosis, while upregulating glutamate signaling genes such as *Gria1*, *Gria4*, and *Slc7a11* ^56–58^ (**Figure 4A, Table S3**). We further examined expression of GFAP and C4b in *hCSF1-Grn**^−/−^*** mice engrafted with KOxMG compared to those receiving WTxMG. We found that both genes were markedly upregulated in the KOxMG group; this increase was absent in the WTxMG group (**Figure 4B**). Loss of *Gria4*, upregulation of *C1q/C4b*, and astrogliosis have previously been shown to drive thalamic hyperexcitability in *Grn^−/−^* mice^8,47^; their normalization following WTxMG xenograft therefore indicates an anti-hyperexcitable effect. To further assess reactive and inflammatory signatures, we examined expression of canonical interferon-stimulated genes, including *Ifi27, Stat1, Stat2, Parp9, Parp14,* and *Trim30*. Consistently, *Grn**^−/−^*** mice exhibited robust upregulation of these interferon-related genes, and restoring progranulin specifically in microglia was sufficient to reduce their expression across the board (**Figure 4C**). These findings demonstrate that microglial progranulin replacement not only dampens complement activation and astrogliosis but also broadly suppresses interferon-driven inflammatory signaling, collectively supporting a shift toward a less reactive and less hyperexcitable thalamic environment. The more robust changes observed in the thalamus are consistent with the well-established sensitivity of this region to progranulin deficiency and its prominent involvement in thalamocortical dysfunction across *Grn*-associated neurodegenerative disorders. Oligodendrocytes (OLs) in the hippocampus were shaped differentially by WTxMG and KOxMG. WTxMG suppressed OL expression of *C4b* and boosted genes for precursor migration (*Robo1*^59^), neural development (*Nrg3*^60^), survival (*Birc2*^61^), and ubiquitin ligase *Trim59* (**Figure S3D, Table S2**). We additionally observed that WTxMG rescued the loss of subcluster 0 seen in KOxMG, while KOxMG selectively expanded subcluster 1 (**Figure S3E**), a population whose transcriptional profile significantly correlated with the disease-associated OL phenotype (**Figure S3F**).

**Figure 4.**
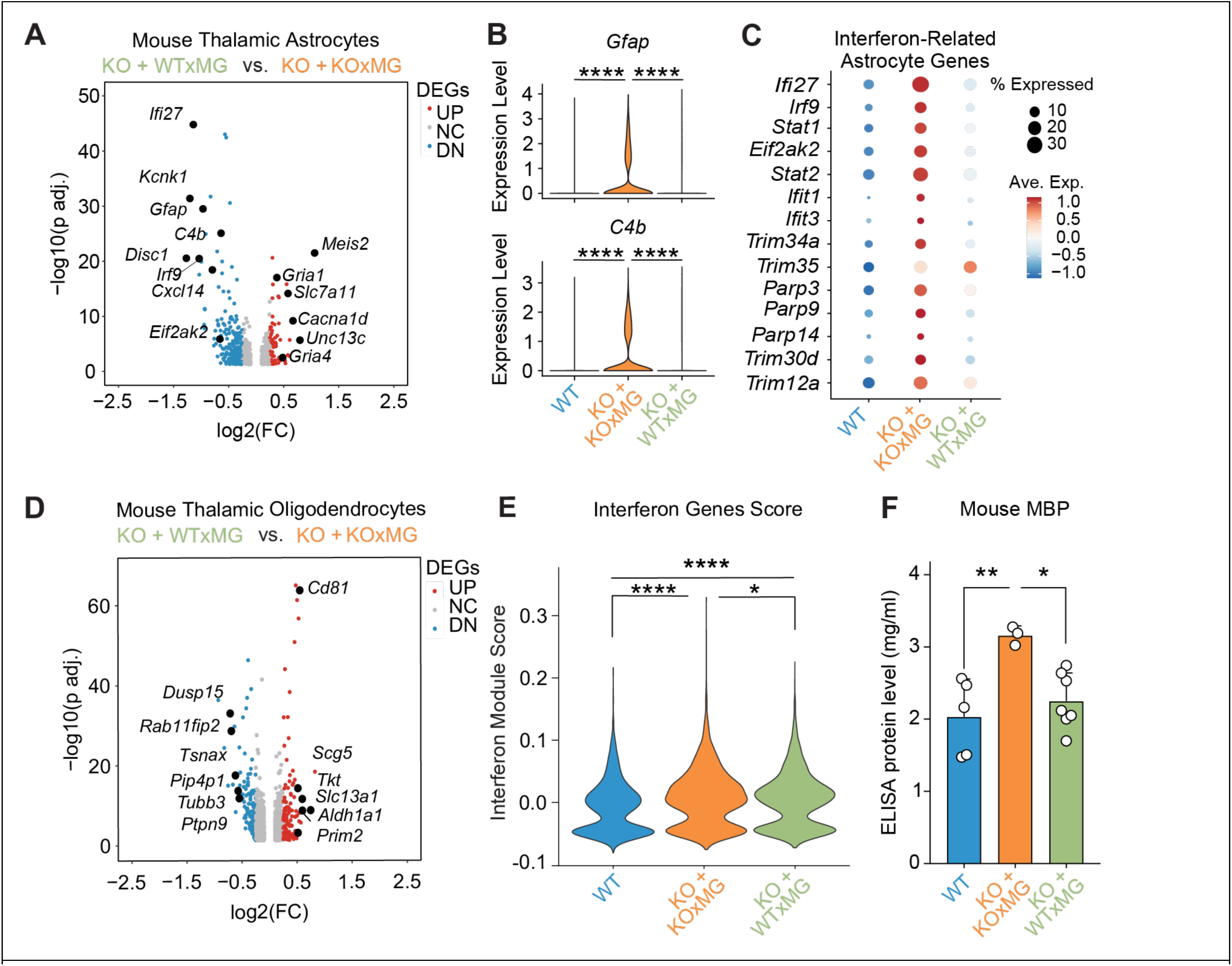
Xenografted human KOxMG or WTxMG exert differential effects on mouse thalamic astrocyte and oligodendrocyte transcriptomes. **(A)** Volcano plot of pseudobulk differentially expressed genes (DEGs) in thalamic astrocytes between WTxMG and KOxMG groups shows significant upregulation and downregulation of multiple genes in the KOxMG group. **(B)** Violin plot for expression of *Gfap* and *C4b* expression in hippocampal astrocytes (\*\*\*\**p* < 0.0001). **(C)** Dotplot of interferon-related genes. **(D)** Volcano plot of pseudobulk DEGs in thalamic oligodendrocytes between WTxMG and KOxMG groups highlights significant transcriptional differences in the KOxMG group. **(E)** Quantification of Interferon Genes Score (\*\*\*\**p* < 0.0001 and \**p* = 0.0129 respectively). **(F)** Quantification of mouse MBP in soluble brain extracts in all experimental groups; *p-*values from one-way ANOVA with Tukey’s multiple comparisons test (*F*_2,12_ = 1.759; \*\**p* = 0.0069 and \**p* = 0.0188 respectively).

In the thalamus, WTxMG normalized OL stress and immune programs. WTxMG downregulated differentiation gene *Dusp15*^62^ while restoring *Slc13a1*, whose loss causes degeneration and seizures^63^ (**Figure 4D, Table S3**). To quantify interferon activation in OLs, we generated a refined module score composed exclusively of canonical interferon-stimulated genes (ISGs) (**Figure 4E**, **Table S3**). This IFN_ISG score revealed a clear gradient of interferon signaling across conditions. KO–KO OLs displayed the highest ISG activation, consistent with an amplified innate immune response in the context of severe pathology. KO–WT OLs showed intermediate ISG induction, indicating partial engagement of interferon pathways despite the presence of wild-type microglia. In contrast, WT OLs exhibited minimal ISG expression, reflecting a baseline, homeostatic state. Together, these data demonstrate that OL interferon signaling is progressively elevated, with KO–KO exhibiting the strongest transcriptional interferon response. Interestingly, KOxMG brains accumulated significantly elevated levels of myelin basic protein (MBP), while WTxMG preserved their levels at WT levels, consistent with effective microglial lysosomal clearance of myelin debris^64^ (**Figure 4F**).

### Xenografted human KOxMG and WTxMG exert differential effects on seizure-related genes in thalamic excitatory and inhibitory neurons in *hCSF1-Grn**^−/−^*** mice

In glutamatergic neurons, WTxMG elevated *Cck*, a neuropeptide that dampens seizure propensity and fine-tunes synaptic output^65,66^, and *Tsnax*, which supports synaptic plasticity and DNA repair^67,68^, while suppressing pro-angiogenic *Vegfa*^39^ and *Nwd1,* which plays a key role in neuronal excitability^69^ (**Figure 5A; Table S4**). In contrast, KOxMG engraftment sharply reduced the sodium-channel subunit encoding genes *Scn1a, Scn2a, Scn3a*, and *Scn8a*^70–76^ (**Figure 5B**), whose loss of function is known to result in epilepsy. Comparing our snRNA-seq findings with FTD patient brains, we found that many of these genes that were diminished in KOxMG transplanted mice were similarly reduced in human progranulin-deficient FTD, whereas WTxMG engraftment preserved these transcript levels (**Figure 5C, Table S5)**.

**Figure 5.**
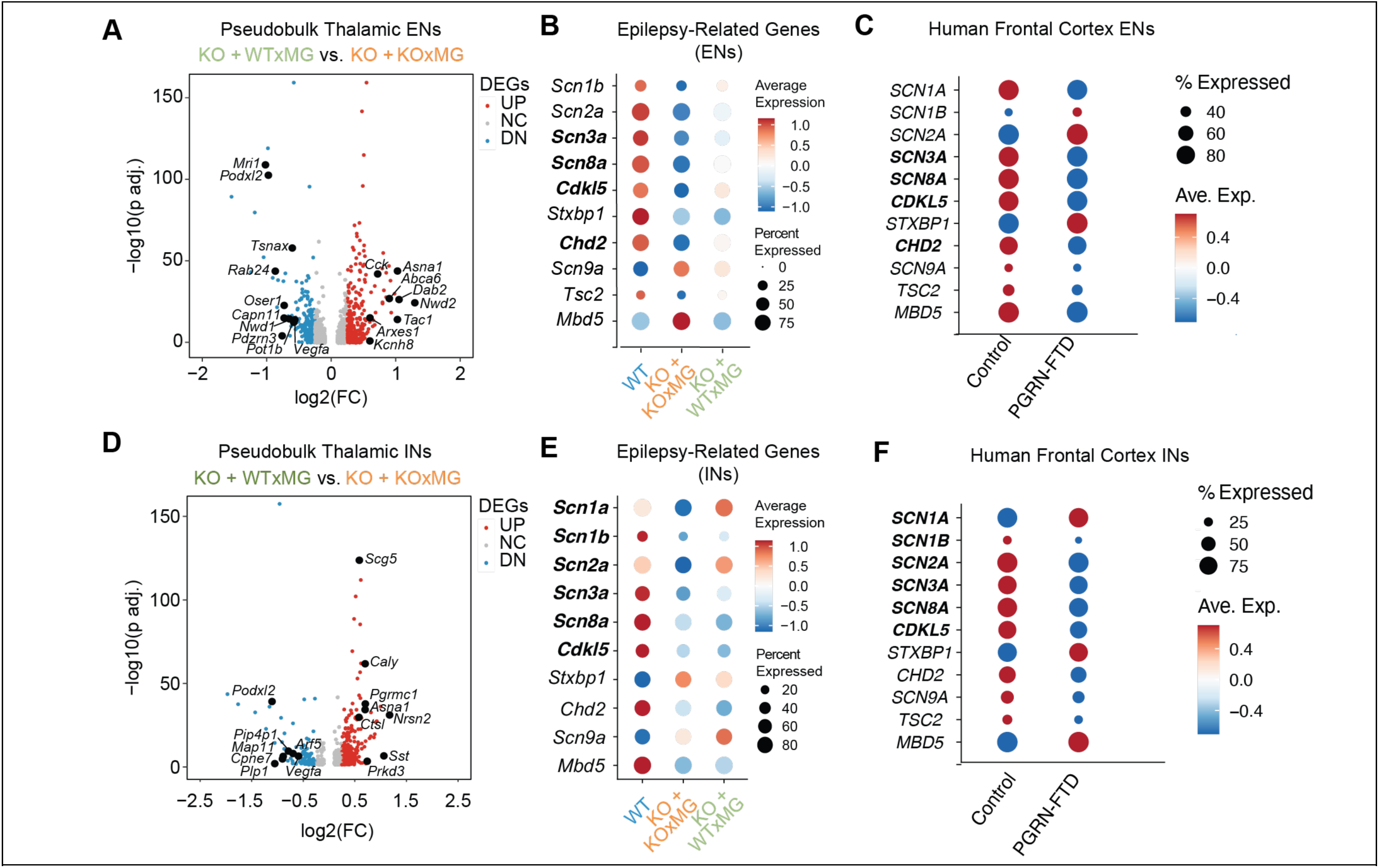
Xenografted human KOxMG and WTxMG exert differential effects on seizure-related genes in thalamic excitatory and inhibitory neurons in *hCSF1-Grn^−/−^* mice. **(A)** Volcano plot of pseudobulk DEGs in thalamic excitatory neurons between WT-xMG and KO- xMG groups, showing upregulation of *Cck, Asna1, Abca6, Dab2, Nwd2, Tac1, Kcnh8, Scube1,* and *Arxes1* and downregulation of *Mri1, Podxl2, Tsnax, Rab24, Nwd1, Oser1, Vegfa, Capn11, Pdzrn3,* and *Pot1b.* **(B)** Dotplot of several epilepsy-related genes, showing reduced expression in the KOxMG group and some restoration in several genes in the WTxMG group (bolded). **(C)** Dotplot of same epilepsy-related genes in human frontal cortex excitatory neurons of control and progranulin-FTD brains, showing a similar pattern as *GRN^+/+^*human xMG (bolded). **(D)** Volcano plot of pseudobulk DEGs in thalamic inhibitory neurons between WTxMG and KOxMG groups, showing upregulation of *Scg5*, *Caly*, *Pgrmc1*, *Asna1*, *Nrsn2*, *Sst*, and *Prkd3* and downregulation of *Podxl2*, *Pip4p1*, *Map11*, *Cpne7*, *Plp1*, *Vegfa*, and *Arf5* in WTxMG injected mice in comparison to those receiving KOxMG. **(E)** Dotplot of several epilepsy-related genes, showing increased expression in the WT group and restoration of several genes in the WTxMG group (bolded). **(F)** Dotplot of the same epilepsy-related genes in human frontal cortex inhibitory neurons of control and Progranulin-FTD brains, showing a similar pattern as *GRN^+/+^* human xMG (bolded).

A parallel pattern emerged in inhibitory neurons: WTxMG upregulated *Caly* and *Nrsn2*, which facilitate receptor endocytosis and vesicle trafficking^77,78^, along with somatostatin, a key modulator of cortical rhythms whose loss marks interneuron vulnerability in AD^79,80^ (**Figure 5D**; **Table S4**). It also suppressed *Vegfa* and the lysosomal protease cathepsin L *(Ctsl*) genes linked to neuroinflammation and synaptic dysfunction^81–83^. KOxMG recipients again exhibited reduced *Scn3a, Scn8a,* and *Cdkl5*, known to result in epilepsy^84^, whereas WTxMG maintained or normalized their levels (**Figure 5E–F; Table S5**). Analysis of human FTD brains compared with control brains showed a similar reduction in many of these epilepsy-related transcripts (**Figure 5F, Table S5**). These results demonstrate the intriguing potential for microglial transplantation to regulate hyperexcitability in *Grn*-deficient brains by modulating critical epilepsy-related gene expression in both excitatory and inhibitory thalamic neurons.

### Human microglia transplantation prevents thalamic hyperexcitability in progranulin-deficient mice

Thalamocortical network dysfunction is thought to underlie multiple behavioral deficits in FTD patients^7,85^. Progranulin deficiency in *Grn^−/−^* mice has previously been shown to induce thalamic network hyperexcitability^85,86^. Having observed the modulation of seizure-related genes by wild-type microglia transplantation in progranulin-deficient mice **(Figure 6)**, we performed multi-unit electrophysiologic recordings^75^. In *ex vivo* thalamic slices **(Figure 6A–C**), analysis of the evoked thalamic spiking to electrical stimulation of the internal capsule axons revealed high-frequency early spiking (within 1.5 s) that stabilized at lower frequencies (delayed, 1.5–3 s) **(Figure 6D–F)**. The frequency of the early spiking response was modestly but significantly increased in *hCSF1-Grn**^−/−^***+KOxMG mice and was normalized in *hCSF1-Grn**^−/−^***+WTxMG mice **(Figure 6G, H)**. However, the frequency of the delayed spiking response was strongly elevated in *hCSF1-Grn-/-*+KOxMG mice but was restored in *hCSF1-Grn**^−/−^***+WTxMG mice to the delayed spiking response observed in *hCSF1* littermates **(Figure 6I, J)**. Taken together, these findings demonstrate that human microglia transplantation prevents the thalamic hyperexcitability observed in *Grn*-deficient mice, underscoring the critical role of microglial function in maintaining thalamic circuit homeostasis and the ability of human WT microglial transplantation to rescue these deficits.

**Figure 6.**
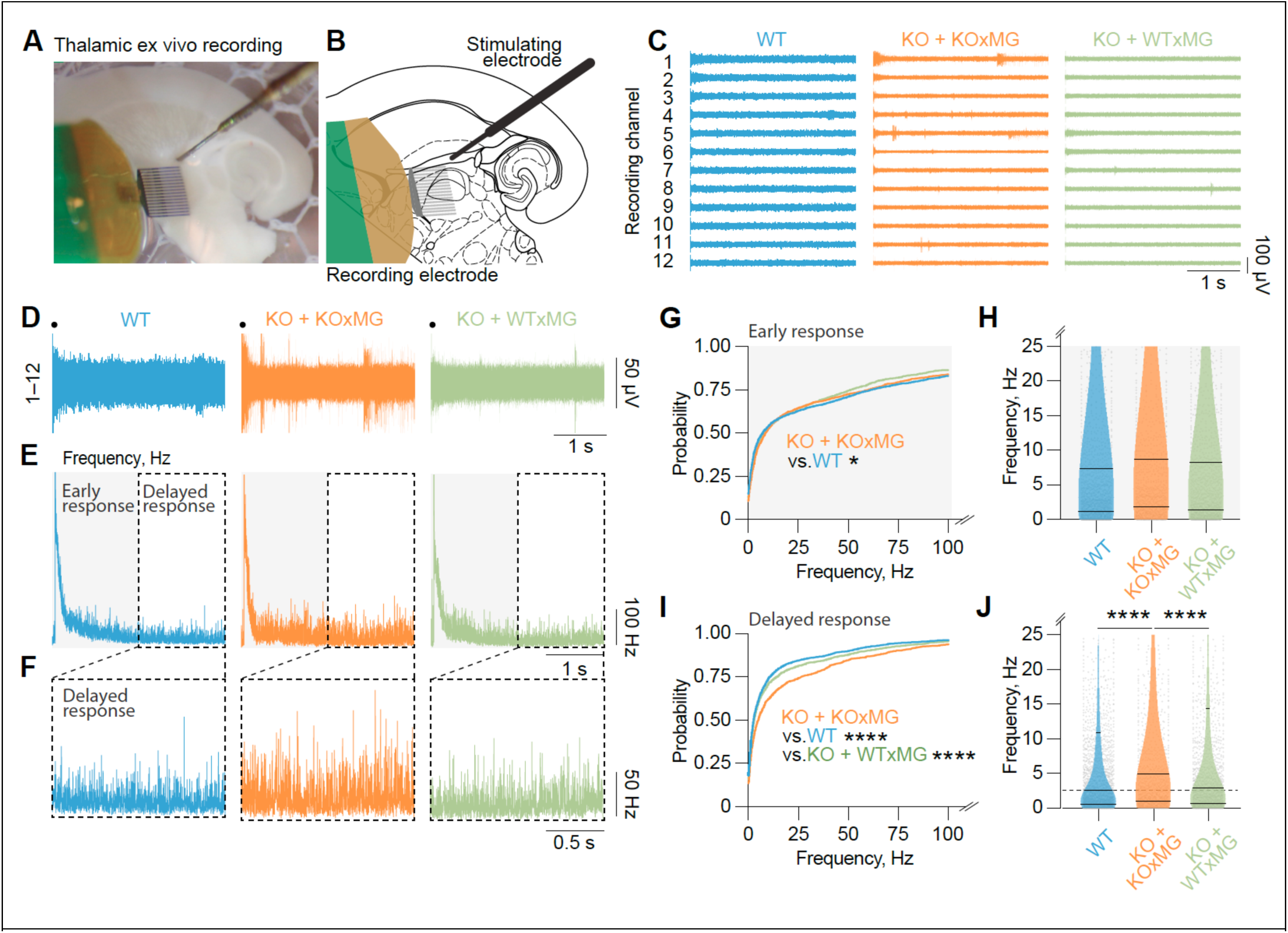
Xenografted human WTxMG rescues thalamic hyperexcitability in *hCSF1-Grn^−/−^* mice. **(A)** Representative image of multi-unit recordings setup of *ex vivo* thalamic horizontal slices in a humidified, oxygenated interface chamber. **(B)** Schematic of multi-unit activity recordings with a Neuronexus 16-channel recording electrode positioned in the thalamus following electrical stimulation of the internal capsule. **(C)** Representative 3-second recording of intrathalamic multi-unit activity evoked by stimulation of the internal capsule (black dot indicates time of stimulation). Only 12 of 16 channels are shown for clarity. **(D)** Collapsed activity from 12 recording channels in (C). Black dot indicates time of stimulation. **(E)** Post-stimulus time histogram of instantaneous spiking frequency from multi-unit activity recorded across 16 channels (all 16 channels were analyzed, 12 representative channels are displayed in (C) from 15 slices from 9 *hCSF1* (WT) mice, 8 slices from 7 *hCSF1-Grn^−/−^* +KOxMG mice, and 12 slices from 9 *hCSF1-Grn^−/−^* +WTxMG mice. The grey box denotes early response (0–1.5 s), and the white box denotes delayed response (1.5–3 s) after stimulation. **(F)** Enlarged instantaneous frequency of the delayed response (1.5–3 s) from (D). **(G)** Relative probability of eliciting spiking during the direct response (0–1.5 s) after stimulation; *p* values from Kolmogorov-Smirnov test with Bonferroni correction (\**p* = 0.0024). **(H)** Instantaneous frequency of spiking during the early response (0–1.5 s) after stimulation; Kruskal–Wallis test, H(3) = 6.767, *p* = 0.0797. **(I)** Relative probability of eliciting spiking during the delayed response (1.5–3 s) after stimulation; *p* values from Kolmogorov-Smirnov test with Bonferroni correction (\*\*\*\**p* < 0.0001). **(J)** Instantaneous frequency of spiking during the delayed response (1.5–3 s) after stimulation; Kruskal-Wallis test, H(3)=56.6, \*\*\*\**p* < 0.0001, and Dunn’s multiple comparisons test (\*\*\*\**p* < 0.0001). Data are presented as mean ± SEM. Comparisons not shown are not significant.

### Human microglia transplantation prevents grooming and social behavioral deficits in *hCSF1-Grn**^−/−^***mice

Hyperexcitability of the thalamocortical circuits has been implicated in OCD-like behaviors, a major clinical presentation in FTD patients^13,87,88^. Consistent with thalamic hyperexcitability, *Grn*-deficient mice are consistently reported to exhibit increased obsessive grooming behaviors^7,17,47^. To assess the therapeutic potential for microglial transplantation to address social behavioral deficits in mice, we analyzed behavioral recordings of transplanted *hCSF1-Grn^−/−^*mice. We assessed grooming behavior in a transparent-bottom open field apparatus (Figure 7A–C). *hCSF1-Grn^−/−^*+WTxMG mice showed reduced obsessive grooming behaviors characterized by repetitive grooming of the snout and body using both front and hind limbs (**Figure 7D–H**). Specifically, the duration of the grooming episodes, but not the frequency, was significantly decreased compared with *hCSF1-Grn^−/−^*+KOxMG mice (**Figure 7D–H**).

**Figure 7.**
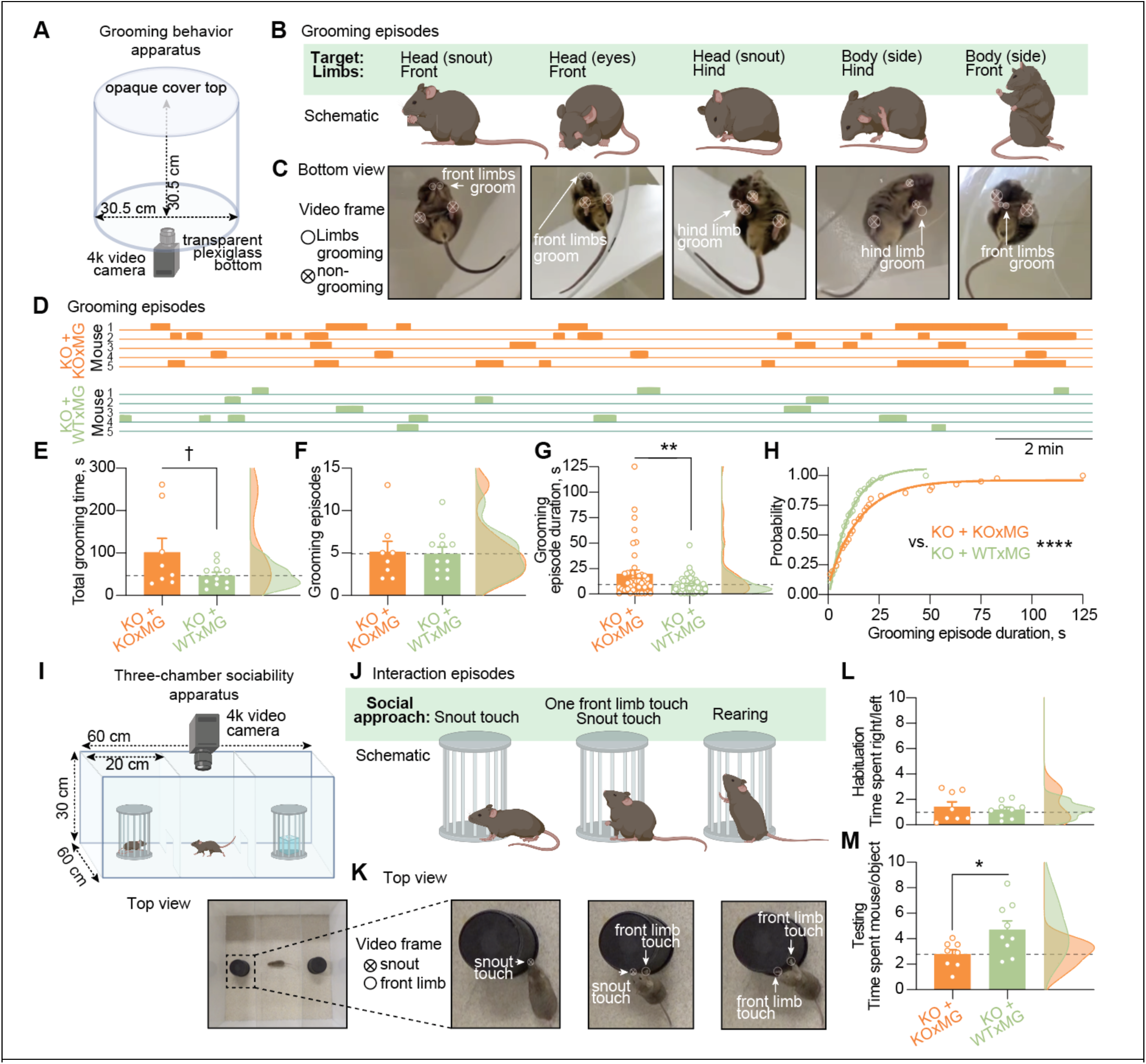
Xenografted human WTxMG rescues grooming and sociability behavioral deficits in progranulin-deficient mice. **(A)** Grooming behavior apparatus consisted of a clear plexiglass cylinder placed on a transparent plexiglass bottom and covered with a white opaque plastic top with a 4k video camera fitted underneath the apparatus. **(B)** Grooming episodes as seen from a side view. Mice groomed both their head and body with front and hind limbs. **(C)** Grooming episodes snapshots as a single video frame corresponding to schematics in B. Open circles denote grooming limbs, crossed—non-grooming. **(D)** Representative episodes of self-grooming during 20-minute recordings in 5 mice from each genotype. **(E)** Quantification of total grooming time, *p* value from Student’s t-test (t(17) = 1.868, ^†^*p* = 0.079). **(F)** Quantification of the number of grooming episodes, *p* value from Student’s t- test (t(17) = 0.1506, *p* = 0.8820). Data represented as mean ± SEM (*hCSF1-Grn^−/−^ +*KOxMG, n = 8; *hCSF1-Grn^−/−^* +WTxMG, n= 11). **(G)** Quantification of the duration of grooming episodes, *p-*value from Student’s t-test (t(93)=2.738, \*\**p* = 0.0074). Data represented as mean ± SEM (*hCSF1-Grn^−/−^ +*KOxMG, n = 41 episodes from 8 mice; *hCSF1-Grn^−/−^ +*WTxMG, n = 54 episodes from 11 mice). **(H)** Probability of grooming episode duration fitted with a nonlinear regression model of exponential plateau curves, *p-*value from extra sum-of-squares F-test (F_3,41_ = 76.74, \*\*\**p* < 0.0001). **(I)** (Top) Three-chamber sociability apparatus comprised of an opaque white acrylic box with two clear plastic inserts with openings with a 4k video camera fitted above the apparatus. (Bottom) Top view of the apparatus with a mouse placed in the middle chamber. **(J)** Interaction episodes of social approach as seen from a side view. Mice approach the pencil wire cup with a 1) snout touch, 2) snout touch and one front limb touch, and 3) rearing (both front limbs touch). **(K)** Interaction episodes snapshots as a single video frame corresponding to schematics in (J). Crossed circles denote snout, open—front limbs. **(L)** Ratio of time spent approaching empty right vs. empty left pencil wire cup during 10 minutes of habituation, *p-* value from Student’s t-test (t(15) = 0.7824, *p* = 0.6335). **(M)** Ratio of time spent approaching the mouse-containing vs object-containing pencil wire cup during 10 minutes of testing, *p* value from Student’s t-test (t(15) = 2.441, \**p* = 0.0275). Data represented as mean ± SEM (*hCSF1-Grn^−/−^ +*KOxMG, n = 8; *hCSF1-Grn^−/−^*+WTxMG, n = 9). \**p* < 0.05, \*\**p* < 0.01. Comparisons not shown are not significant.

Social behavior deficits in FTD patients are characterized by social withdrawal, disinhibition, and loss of sympathy and empathy^89^. *Grn*-deficient mice have also been shown to develop social behavior deficits, most prominently observed in the three-chamber sociability assessment^89–91^. Behavioral analysis of transplanted *hCSF1- Grn^−/−^* mice revealed that animals receiving *GRN*-deficient human microglia (KOxMG) preferentially interacted with an inanimate object rather than a conspecific. In contrast, *hCSF1⁻ Grn^−/−^* mice transplanted with wildtype human microglia (WTxMG) showed an increased preference for social interaction, spending more time engaging with a mouse than with an inanimate object. (**Figure 7I–M**). These findings demonstrate the important role microglia play in regulating social behavior and further highlight the therapeutic potential for microglial transplantation to not only prevent progranulin- FTD-related neuropathologies but to also to rescue behavioral deficits in *hCSF1-Grn^−/−^* mice.

## Discussion

In this study, we sought to dissect the biology of restoring progranulin specifically in microglia using human iPSC-derived microglia transplanted into a xenotolerant, progranulin-deficient FTD mouse model. A single injection of WTxMG, but not KO xMG, reinstated brain-wide progranulin and shifted microglial states toward a more homeostatic profile, providing broad protection against multiple phenotypes associated with progranulin loss. Notably, although both neurons and microglia express progranulin, our findings highlight that microglial progranulin plays a disproportionately important role in shaping neuroimmune homeostasis, neuronal vulnerability, and circuit integrity in the context of FTD. These results emphasize the biological importance of progranulin within microglia and illustrate how microglial progranulin insufficiency propagates dysfunction across many neural cell types.

Our single nuclei analyses further demonstrate that microglial progranulin can strongly influence whether microglia adopt a stable, homeostatic identity or transition into a reactive, inflammatory state. Wild-type human microglia preferentially occupied a homeostatic, anti-inflammatory subcluster. This subcluster was marked by preserved identity genes and downregulation of interferon-related mediators such as STAT1 and IFI44L. In contrast, *GRN*-deficient human microglia disproportionately populated a stress-associated subcluster enriched for activation markers including *IL1RAP*, *TNFAIP2*, *SCN9A*, and *DAPK1*. The strong anticorrelation between these two subclusters reveals that progranulin sufficiency stabilizes a protective transcriptional program while actively suppressing maladaptive inflammatory signaling.

Importantly, the consequences of microglial progranulin restoration extended well beyond the transplanted human cells. Endogenous mouse microglia in *hCSF1-Grn^−/−^* + WTxMG brains exhibited downregulated inflammatory pathways, reduced the abundance of disease-associated subclusters, and shifted transcriptional profiles consistent with a homeostatic profile. Conversely, microglia in *hCSF1-Grn^−/−^* +KOxMG mice showed elevated inflammatory gene expression, including induction of *Cst3*, *Hexb*, *Ly6e*, and other innate immune regulators. These findings reveal a powerful non–cell-autonomous effect in which progranulin-sufficient microglia broadcast cues that remodel the molecular states of neighboring microglia across the brain.

Restoring progranulin specifically in microglia produced widespread, cell-non-autonomous effects. These effects extended to surrounding mouse glia and neurons, which depend on normal microglial signaling for homeostasis. Even at physiologic levels, microglial progranulin was sufficient to suppress the aberrant activation of complement and interferon pathways, which were strongly upregulated in *hCSF1-Grn^−/−^*astrocytes and oligodendrocytes. This normalization is consistent with progranulin’s known role in restraining lysosomal stress, dampening inflammatory signaling, and limiting microglial release of cytokines and complement components that secondarily activate neighboring glia^4,5,92^. In neurons, both excitatory and inhibitory, microglial progranulin restored transcriptional signatures disrupted by progranulin deficiency, notably reversing the aberrant upregulation of epilepsy-associated genes that are typically induced by heightened complement signaling, synaptic pruning, and inflammatory cytokines^93–96^. Notably, the neuronal gene expression profiles in mice xenografted with wild-type human microglia shifted toward those of control human brains, indicating that microglial progranulin regulates neuronal homeostasis through conserved pathways that modulate synaptic maintenance, network excitability, and stress-response programs^24,47,97^.

At the functional level, these transcriptional improvements translated into measurable rescue of circuit activity. Reintroducing progranulin in microglia alone was sufficient to prevent thalamic hyperexcitability, a phenotype in progranulin deficiency driven by excess complement deposition, heightened microglial reactivity, altered astrocytic buffering, and reduced inhibitory tone. Restored microglial progranulin likely acts through multiple converging mechanisms, including normalization of lysosomal function, suppression of microglial C1q and cytokine production, and stabilization of synaptic integrity, to preserve neuronal firing properties and prevent runaway excitation^92,98–101^. Behaviorally, mice engrafted with wild-type human microglia also showed significant reductions in total grooming time and the duration of individual grooming bouts, consistent with reduced network hyperexcitability and improved corticostriatal regulation. Together, these findings demonstrate that microglial progranulin is a central regulator of glial–neuronal cross-talk, synaptic stability, and circuit homeostasis, and position microglia as a powerful therapeutic target for neurodegenerative diseases characterized by excitability and network dysfunction.

The minimal transcriptional differences observed between KO–WT and KO–KO endogenous mouse microglia indicate that these cells are unlikely to be the primary drivers of the widespread changes detected in surrounding glial and neuronal populations. Instead, our data strongly support a model in which restoration of microglial progranulin is sufficient to reshape the local cellular environment, with downstream effects on astrocytes, oligodendrocytes, and neurons occurring independently of major host microglial reprogramming. While the endogenous mouse microglia of the KO–WT group do exhibit shifts toward a less inflammatory state, these limited changes cannot account for the magnitude of non-cell-autonomous effects observed across other cell types, reinforcing transplanted human microglia as the principal source of progranulin-dependent circuit remodeling.

Although progranulin is classically recognized for its lysosomal functions, recent work suggests that granulins may carry distinct biological activities^8^. This raises the intriguing possibility that microglial processing of progranulin, rather than absolute protein levels alone, contributes to establishing the homeostatic and anti-inflammatory programs we observe. Our data underscore that proper handling of progranulin within microglia is critical for maintaining immune quiescence, synaptic stability, and broader neural circuit homeostasis^14,15,47,92,102^.

While our findings provide clear insight into how microglial progranulin shapes cellular and network states, several questions remain. One consideration is that the homozygous *Grn**^−/−^*** model represents complete progranulin loss, which more closely resembles neuronal ceroid lipofuscinosis (NCL) than the haploinsufficiency characteristic of GRN-FTD. However, this stringent model provides a robust test of microglial progranulin sufficiency, and these findings may have direct therapeutic relevance for patients with NCL harboring biallelic GRN mutations. Our xenograft model captures key aspects of human microglial biology, but does not fully resolve how progranulin-dependent microglial states evolve across all brain regions or over extended timescales. In addition, the molecular intermediates through which progranulin exerts its effects, whether the full-length protein, granulins, or downstream lysosomal signals, remain to be defined. Future studies aimed at dissecting these pathways and tracking microglia–neuron interactions *in vivo* will help clarify the mechanisms by which microglial progranulin maintains immune homeostasis and supports neural circuit stability. Overall, our work establishes microglial progranulin as a key determinant of cellular resilience and coordinated network function in the progranulin-deficient FTD brain.

## MATERIAL AND METHODS

### Animals

All animal procedures were conducted in accordance with guidelines set forth by the National Institutes of Health and the University of California, Irvine Institutional Animal Care and Use Committee, and the University of California, San Francisco Institutional Animal Care and Use Committee. *hCSF1-Grn^−/−^* mice (abbreviated as *hCSF- Grn^−/−^*) were generated by backcrossing (*Grn-/-* mice, Jackson Laboratories #029919) with xenotransplantation compatible hCSF1 mice (Jackson Laboratories #017708), harboring deletions in *Rag2* and *Il2rg* and expressing humanized CSF-1. The resulting *hCSF1-Grn^−/−^*and strain-matched *hCSF1-Grn^+/+^* littermate controls were maintained on a mixed BALB/c × B6.129 background. All mice were age-matched and group-housed on a 12 h:12 h light/dark cycle with ad libitum access to food and water. Mice were housed under standard ambient temperature and humidity conditions. Equal number of male and female mice were used for experiments.

### Induced pluripotent stem cell lines

The parental WTC11-mEGFP (AICS-0036-006, Coriell) human iPSC line^20,95,96^ was acquired through the Allen Cell Collection (Coriell Institute for Medical Research), and used in the first study (**Figure 1**). Isogenic *GRN^−^/^−^* human iPSCs were generated on a WTC11 iPSC background. WTC11 is a widely used and previously validated iPSC line derived from a healthy 30-year-old male (Coriell # GM25256)^97,98^. Two sgRNAs were designed to delete a 5231 bp fragment within the progranulin locus. One sgRNA targeted the upstream region of the *GRN* promoter (AGCTCAAGGAGATGCTCCTA), while the other specifically recognized the downstream region of the exon containing the ATG start codon of the *GRN* gene (GCCAATCCAAGATGACCCTT). iPSCs were collected following Accutase enzymatic digestion and resuspended in human stem cell nucleofector buffer (Lonza). Ribonucleoprotein (RNP) complex was generated by incubating HiFi Cas9 Nuclease (IDTDNA) with guide RNAs for 15 min at 23 °C. The cell suspension was then nucleofected with 50 μg of RNP complex using Amaxa Nucleofector program B-016. To screen for the deletion, primers flanking the regions outside of the two sgRNA target sequences were designed, and positive clones were identified by PCR. Positive clones were further characterized using PCR with three primers: two outside the sgRNA target sequences and one within the deleted region. Monoallelic targeting was identified by the appearance of two distinct products—one of wild-type size and one of targeted size. The PCR products from homozygous clones were validated by Sanger sequencing to confirm the deletion and the integrity of the locus following CRISPR-Cas9 RNP editing. Homozygous clones were then expanded and re-characterized by PCR and Sanger sequencing. All cells were verified to be karyotypically normal and tested negative for mycoplasma contamination.

### Differentiation of hematopoietic progenitor cells from human iPSCs

Hematopoietic progenitor cells (HPCs) were differentiated following the highly replicated protocol published by McQuade et al^95^. To begin HPC differentiation, WT and KO iPSCs were passaged onto 6-well Matrigel-coated plates in TeSR-E8 at a density of 80 colonies per 35-mm well, per 35-mm well, with approximately 100 cells per colony. On day 0, cells were transferred to Medium A from the STEMdiff Hematopoietic Kit (05310; STEMCELL Technologies). On day 3, cells were exposed to Medium B and maintained in Medium B for 7 additional days while small round HPCs began to lift off the colonies. On day 10, non-adherent CD43^+^ HPCs were collected by carefully removing medium and non-adherent cells with a serological pipette. At this point, HPCs were frozen in BamBanker freezing solution (Wako, NC9582225) for long-term storage. Cells used for transplantation were later thawed in complete iMG medium for 18–24 h to allow recovery, before being resuspended at 62,500 cells/µl in 1× DPBS (low Ca^2+^, low Mg^2+^) for transplantation. A total of 500,000 HPCs were injected per mouse.

### Experimental design

In the first study, *hCSF1-Grn^−/−^* mice were transplanted with WTC11-GFP HPC (WTxMG) at P2. Control groups (*hCSF1-Grn^+/+^* and *hCSF1-Grn^−/−^*mice) were injected with the same volume of 1xPBS. Mice were terminated at 12 months of age, and biochemical and histological analyses were performed. In the second study, *hCSF1-Grn^−/−^*mice (KO) were transplanted either with WTxMG or KOxMG at P2, and, as a control group, WT mice did not receive any treatment. Behavioral tests were performed at 9.5 months of age, and electrophysiological assays were performed at 10.5–11.5 months of age in these same mice. This sequential design ensured that behavioral phenotypes were established prior to terminal procedures, minimizing handling stress that might confound molecular and circuit-level analyses. Mice were euthanized and perfusion. Immunohistochemistry, RNA sequencing, and biochemical analysis were performed with collected brain samples. The two studies employed different control strategies suited to their respective aims. In the first study, PBS-injected *hCSF1-Grn^−/−^*mice served as controls to confirm that the injection procedure itself did not account for observed rescue effects. In the second study, KO mice transplanted with *GRN*-deficient microglia (KOxMG) served as the disease control, as these cells lack the capacity to restore progranulin. The primary comparison was therefore between WTxMG and KOxMG, directly testing whether microglial progranulin is required for phenotypic rescue.

### Intracranial Transplantation at post-natal days 1–2

All mouse surgeries and use were performed in strict accordance with approved National Institutes of Health (NIH) and institutional guidelines. HPCs were transplanted into P1–2 *hCSF1-Grn^−/−^*mouse pups via intracerebroventricular injection as previously described^95^. Using a 10μL Hamilton syringe, cryoanesthetized mice received 2 μl per injection, for a total of 4 injections, of HPCs suspended in sterile 1X PBS at 125,000 cells/μl per injection site. Bilateral injections were performed at two-fifths of the distance from the lambda suture to each eye, injecting into the ventricles at 3 mm depth and into the overlying cortex at 1 mm depth. After injection, the pups were allowed to recover on a warming Deltaphase pad with a cloth covering to prevent rapid rewarming damage. wrapped around the warming pad to prevent rapid rewarming damage and then transferred back to the mother’s cage. Control groups (*hCSF1-Grn^+/+^* and *hCSF1-Grn^−/−^*mice) were injected with saline (vehicle control) as described above.

### Brain collection and isolation of Soluble Protein Homogenates

Animals were perfused (20 mL/min) with 1X DPBS (4 °C) for 3 min. Brains were surgically excised from each mouse and cut in half along the mid-sagittal plane. The left hemisphere was placed in 4% (wt/vol) paraformaldehyde (PFA) for 36 h at 4 °C for subsequent immunohistochemistry (IHC), and the right hemisphere was fresh-frozen on dry ice and stored at -80 °C for biochemical analysis and proteomics. Fresh-frozen hemispheres were crushed in liquid nitrogen using mortar and pestle. The brain powder was homogenized in solution of T-PER (78510, Thermo Scientific, Waltham, MA) and phosphatase and protease inhibitor mixtures (78426 and 78426, Thermo Scientific, Waltham, MA). Homogenates were centrifuged at 16,000 rcf (relative centrifugal force) for 30 min at 4 °C, and supernatants were stored at −80 °C for analysis. Protein concentration of soluble brain samples was determined using the BCA Protein Assay Kit (Pierce), and 2 mg/mL of total protein for each sample was run for ELISAs.

### Quantitative Biochemical Analysis

Analysis of soluble brain samples was conducted using commercially available ELISAs following the manufacturer guidelines to measure human Progranulin (DY2420; R&D Systems), mouse Myelin Basic Protein (MBP; ABIN6957934; antibodies-online.com), and V-PLEX Plus Mouse Cytokine 19-Plex Kit (K15255G; Meso Scale Discovery, MSD). Protein levels for each sample were then calculated via comparison to the assay-specific standard curve.

### Immunohistochemistry

Before sectioning, hemibrains fixed with 4% PFA were cryoprotected in a 30% (wt/vol) sucrose solution at 4 °C. Brains were sectioned coronally into 30-μm-thick sections on a freezing microtome (Leica, SM 2010R) and stored in a solution of 0.05% NaN3 (S2002; Sigma-Aldrich) in 1× PBS (P44017-100TAB; Sigma-Aldrich) as free-floating slices. For staining, tissue was blocked for 1 h in 1× PBS, 0.2% Triton X-100 (9002-93-1; Thermo Fisher Scientific), and 10% donkey serum (NC9624464; ThermoFisher Scientific). Immediately following blocking, brain sections were placed in primary antibodies diluted in 1× PBS and 1% donkey serum and incubated overnight with gentle shaking at 4 °C. Samples were then incubated in conjugated secondary antibodies for 1 h, followed by mounting on microscope slides. Sections were labeled with combinations of goat anti-IBA1 (1:300; ab5076; Abcam), rabbit anti-IBA1 (1:300; 019-19741; Wako), rabbit anti-Ku80 (1:200; ab80592; Abcam), rat anti-Lamp1 (1:300; ab25245; Abcam), chicken anti-GFP (1:500; AB16901; Millipore-Sigma), and rat anti-CD68 (1:200; MCA1957; Bio-Rad) antibodies overnight at 4 °C followed by 1 h incubation at 23 °C with gentle shaking. After washing with 1× PBS for 5 minutes three times, sections were incubated with highly cross-adsorbed AlexaFluor-conjugated secondary antibodies (1:400; ThermoFisher) for 1 h in the dark, then washed three times with 1× PBS before mounting with Fluoromount-G (0100-01; SouthernBiotech).

### Imaging acquisition and processing

Immunofluorescent sections were visualized and captured using an Olympus FX3000 confocal microscope using identical confocal and Z-stack settings for a given quantitative comparison. High-resolution hemibrain stitching was performed using Fluoview FV31S-DT software. To calculate autofluorescence and CD68 immunoreactivity, three consecutive sections were imaged per region per animal, then analyzed using the “Surfaces” function in IMARIS software to report a single value per mouse (Bitplane). All images were examined blinded to genotype and treatment using batch processing. For some representative images, brightness and contrast settings were slightly and equally adjusted across groups to reveal fine structures and morphology. Importantly, no such changes were made to any images used for quantification.

### Single-nuclei RNA Sequencing

Nuclei isolation from frozen mouse hippocampi was adapted from a previous study^99^, with modifications. All procedures were performed on ice or at 4 °C. In brief, postmortem brain tissue was placed in 1,500 µl of Sigma nuclei PURE lysis buffer (Sigma, NUC201-1KT) and homogenized with a Dounce tissue grinder (Sigma, D8938-1SET) with 15 strokes with pestle A and 15 strokes with pestle B. The homogenized tissue was filtered through a 35-µm cell strainer, centrifuged at 600g for 5 min at 4 °C and washed three times with 1 ml of PBS containing 1% bovine serum albumin (BSA, Thermo Fisher Scientific, 37525), 20 mM DTT (Thermo Fisher Scientific, 426380500), and 0.2 U/µl recombinant RNase inhibitor (Ambion, AM2684). Nuclei were then centrifuged at 600g for 5 min at 4 °C and resuspended in 350 µl of PBS containing 0.04% BSA and 1× DAPI, followed by fluorescence-activated cell sorting to remove cell debris. The sorted suspension of DAPI-stained nuclei was counted and diluted to a concentration of 1,000 nuclei per µl in PBS containing 0.04% BSA. For droplet-based snRNA-seq, libraries were prepared with Chromium Next GEM Single Cell 3′ Kit v3.1 (10x Genomics, PN-1000268) according to the manufacturer’s protocol (CG00315 Rev E). The snRNA-seq libraries were sequenced on a NovaSeq 6000 sequencer (Illumina) with 100 cycles. Gene counts were obtained by aligning reads to the GRCh38 and mm10 dual references with the Cell Ranger software (v.6.1.2; 10x Genomics). To account for unspliced nuclear transcripts, reads mapping to pre-mRNA were counted. Cell Ranger v6.1.2 default parameters were used to call cell barcodes. Human or mouse species identity was assigned with 80% of UMIs mapped uniquely to that species. We further removed genes expressed in fewer than three cells and removed cells with fewer than 100 unique genes. Normalization and clustering were done with the Seurat package v4.0.0 ^100^. In brief, the raw RNA counts were transformed using Seurat function SCTransform. Principal components analysis (PCA) was performed using the normalized SCT dataset (RunPCA function), and two-dimensional representations were generated using the top 30 principal components as input to the RunUMAP function. Cell clusters were identified with the Seurat functions FindNeighbors (using the top 15 principal components) and FindClusters. To mitigate potential batch effects, sample integration was performed using FindIntegrationAnchors and IntegrateData functions in Seurat. For each cluster, we assigned a cell-type label using statistical enrichment for sets of marker genes and manual evaluation of gene expression for small sets of known marker genes. Differential gene expression analysis was done using the FindMarkers function and MAST^103^. Genes were considered significantly differentially expressed at an FDR-adjusted *p*-value < 0.05 and |log2 fold change| > 0.1. To identify gene ontology and pathways enriched in the DEGs, DEGs were analyzed using the MSigDB gene annotation database^104,105^. To control for multiple testing, we used the Benjamini–Hochberg approach to constrain the FDR.

### Human FTD Patient Data Single-nuclei RNA Sequencing

Patient information for single-nuclei RNA sequencing (sn-RNAseq) analysis performed on human FTD patients (n = 8) and control patients (n = 7) can be found in the first tab of **Table S5**. Human FTD snRNA-seq data is available at the GEO archive under accession number GSE250280, and specific methods about tissue acquisition, processing, and analysis have been previously described^103^.

### Electrophysiology

Slice preparation was performed as previously described^64^. Briefly, mice were euthanized with 4% isoflurane, then transcardially perfused with ice-cold cutting solution containing 234 mM sucrose, 2.5 mM KCl, 1.25 mM NaH2PO4, 10 mM MgSO4, 0.5 mM CaCl2, 26 mM NaHCO3, and 11 mM glucose, equilibrated with 95% O2 and 5% CO2, pH 7.4. Following perfusion, mice were rapidly decapitated, and brains were immediately immersed in the ice-cold sucrose cutting solution. Thalamic slice preparation was performed as described ^8,64,80^. Horizontal thalamic slices containing the somatosensory ventrobasal (VB) thalamus and the nucleus reticularis thalami (nRT) were cut at 400 μm (for thalamic microcircuit studies) with a Leica VT1200 microtome (Leica Microsystems). Slices were incubated at 32 °C for 30 min, then at 24–26 °C for 1 hour, in artificial cerebrospinal fluid (aCSF) containing 126 mM NaCl, 2.5 mM KCl, 1.25 mM NaH2PO4, 1 mM MgCl2, 2 mM CaCl2, 26 mM NaHCO3, and 10 mM glucose, 300–310 mOsm, equilibrated with 95% O2 and 5% CO2, pH 7.4.

#### Recordings

Horizontal slices (400 μm), known to preserve the connectivity between the somatosensory VB and nRT thalami, were placed in an interface chamber at 34 °C and superfused at a rate of 2 mL/min with oxygenated aCSF supplemented with 0.3 mM glutamine for cellular metabolic support and 5 nM apamin to increase thalamic microcircuit evoked responses. Extracellular multi-unit activity (MUA) recordings were obtained with a linear 16-channel multi-electrode array (Neuronexus) that spanned the nRT and VB thalamic regions. MUA signals were amplified 10,000 times and band-pass filtered between 100 Hz and 6 kHz using the RZ5 system (Tucker-Davis Technologies,TDT, SCR_006495). The position of the recording array was visually verified for each recording to confirm the position of electrodes in the thalamic subregions. Electrical stimuli were delivered to the internal capsule with bipolar tungsten microelectrodes (50–100 kΩ, FHC). The stimuli were 100 μs in duration, 50 V in amplitude, and delivered every 30 s for 10–20 trials.

**Data analysis** was performed offline after the recordings were complete. Extracellular spikes from multi-unit activity (MUA) were detected with custom MATLAB scripts (Huguenard Lab, Stanford University) by taking the first derivative of the signal and thresholding relative to baseline root mean square (RMS) values. Spikes with waveform durations longer than 2 ms were excluded. Parameter settings were confirmed by an experienced experimenter to optimize spike detection across recordings. For automatic spike detection, the first three seconds of each trial were analyzed. The evoked activity occurring 0–1.5 s after the electrical stimulation of each trial was defined as the early response, whereas activity occurring 1.5–3 s after stimulation was defined as the delayed response. Extracellular spiking frequency was calculated by pooling spikes over 1 ms bins, averaging across the 16 channels, and then averaging across slices. Each mouse had 1–3 slices recorded. Individual slices were treated as independent observations, as each slice represents a distinct preparation with independent handling and recording conditions. Both slice and animal numbers are reported in figure legends. For statistical analysis, relative spiking probabilities during the early and delayed responses was downsampled 10-fold and compared using the Kolmogorov–Smirnov test with Bonferroni correction for multiple comparisons. Absolute instantaneous spike frequencies were analyzed using the Kruskal–Wallis test with Dunn’s correction for multiple comparisons.

### Behavior

#### Grooming

To quantify grooming behaviors, mice were transferred from the vivarium into a quiet room for an hour prior to behavioral testing. Each mouse was placed in an open field circular arena cylinder (30.5 cm in diameter) and video recorded at 30 frames/s for 20 min using a camera positioned 12 inches below the cylinder. The video-recorded sessions were scored for grooming behaviors involving the front and hind paws, including grooming of the head (snout, eyes) and body (neck, back, sides and hind paws) by a trained observer blinded to genotype and treatment, at a 0.25x speed of the original recording. The onset time and duration of each grooming episode were recorded. Total grooming time, the number of grooming episodes, and grooming episode durations were analyzed using Student’s t-test. The probability of grooming episode duration was fitted with a nonlinear regression model of the exponential plateau and analyzed statistically with an extra sum-of-squares F test.

#### Three-chamber sociability test

The three-chamber sociability test was adapted from previously described methods^79,101^. The apparatus was an opaque white acrylic box with a tan bottom (60 cm × 60 cm) and 30-cm-high walls. Two transparent acrylic insert dividers with 5 cm × 5 cm openings were used to create three chambers (each 60 cm × 20 cm). A 4K Logitech video webcam was placed 12 inches above the apparatus. For habituation, mice were placed in the middle chamber and allowed to freely explore the apparatus with two empty inverted pencil wire cups placed in the right and left chambers for 10 minutes. For testing, mice were directed to the middle chamber, the openings to the side chambers were closed, and a Lego block object and a mouse (adult sex-matched C57BL/6J) were placed into the inverted pencil wire cups in the side chambers. The location of the object and mouse placements were alternated equally during the experiment. After opening the side chambers, mice were allowed to explore the Lego block object or mouse through the pencil wire cups for 10 minutes. Active approach was scored by a blinded observer as: 1) snout touch, 2) snout touch plus one front limb touch, 3) rearing (both front limbs touching). Active approach time spent with right/left pencil wire cups and time spent with pencil wire cups containing mouse/object were analyzed using Student’s t-test after confirming normality by Shapiro-Wilk tst.

### Statistical Analyses

GraphPad Prism 9 was used to perform statistical tests and generate P values. We used standard designation of P values throughout the Figures (ns = not significant (p ≥ 0.05); \**p* < 0.05; \*\**p* < 0.01; \*\*\**p* < 0.001; ****p < 0.0001). Values are represented as mean ± SEM. Details of the number of replicates and the specific statistical test used are provided in the individual figure legends. Data for all antibodies were calculated from an average of three matched coronal sections per animal.

## LIST OF SUPPLEMENTARY MATERIALS

**Table S1:** List of marker genes, DEGs, and upstream pathway analyses for xenotransplanted human microglia. Related to **Figure 3**.

**Table S2:** List of marker genes and DEGs for hippocampal mouse astrocytes and oligodendrocytes. Related to **Figure 4**.

**Table S3:** List of marker genes and DEGs for thalamic mouse astrocytes and oligodendrocytes. Related to **Figure 4**.

**Table S4**: List of marker genes and DEGs for thalamic mouse excitatory neurons and inhibitory neurons. Related to **Figure 5**.

**Table S5:** Human control and FTD patient information, and lists of marker genes and DEGs for human patient excitatory neurons and inhibitory neurons. Related to **Figure 5**.

**Figure S1.**
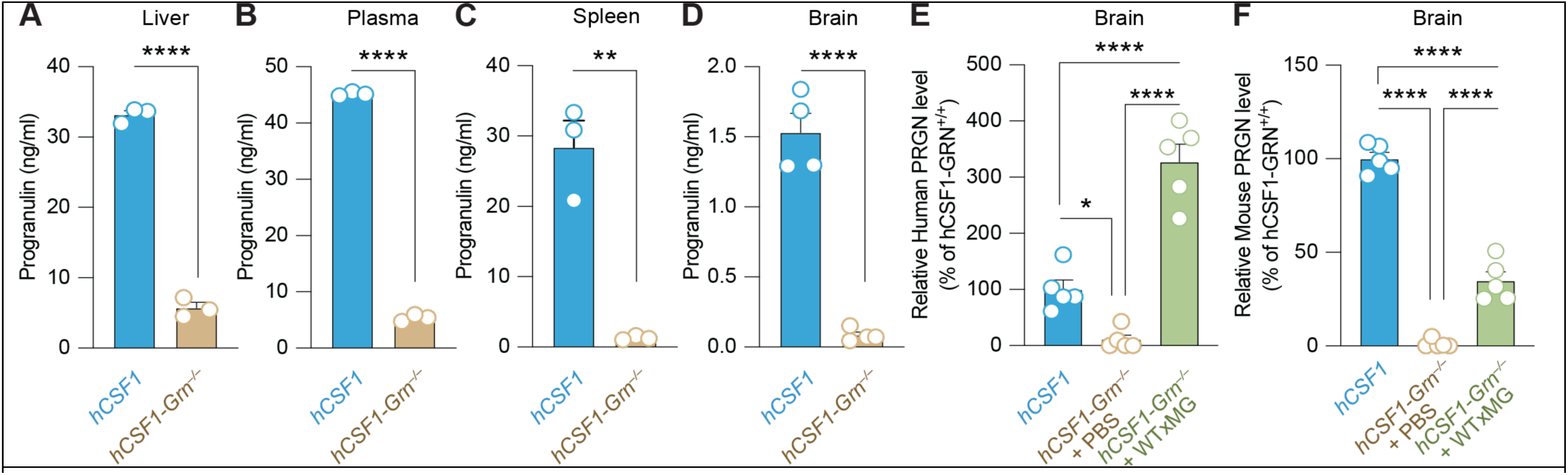
Progranulin protein levels within peripheral organs and the brain. Levels of murine progranulin within the liver (t=27.60, \*\*\*\**p* < 0.0001) **(A)**, plasma (t=96.25, ****p < 0.0001) **(B)**, and spleen (t=7.07, \*\**p* = 0.0021) **(C)** of *hCSF1-WT* and *hCSF1-Grn^−/−^*littermates were measured by ELISA. N = 3 mice/group, all unpaired two-tailed t-test. **(D)** Progranulin levels measured within the brain (t=10.31, \*\*\*\**p* < 0.0001). N = 4–5 mice/group, respectively, unpaired two-tailed t-test. **(E)** Progranulin levels measured within the brain using a human-targeted ELISA, n = 5 samples/group, one-way ANOVA with Tukey’s multiple comparisons test (F_2,12_=1.803, \*\*\*\**p <* 0.0001). **(F)** Progranulin levels measured within the brain using a mouse-targeted ELISA, n = 5 samples/group, one-way ANOVA with Tukey’s multiple comparisons test (F_2,12_=2.80, \*\*\*\**p <*0.0001).

**Figure S2.**
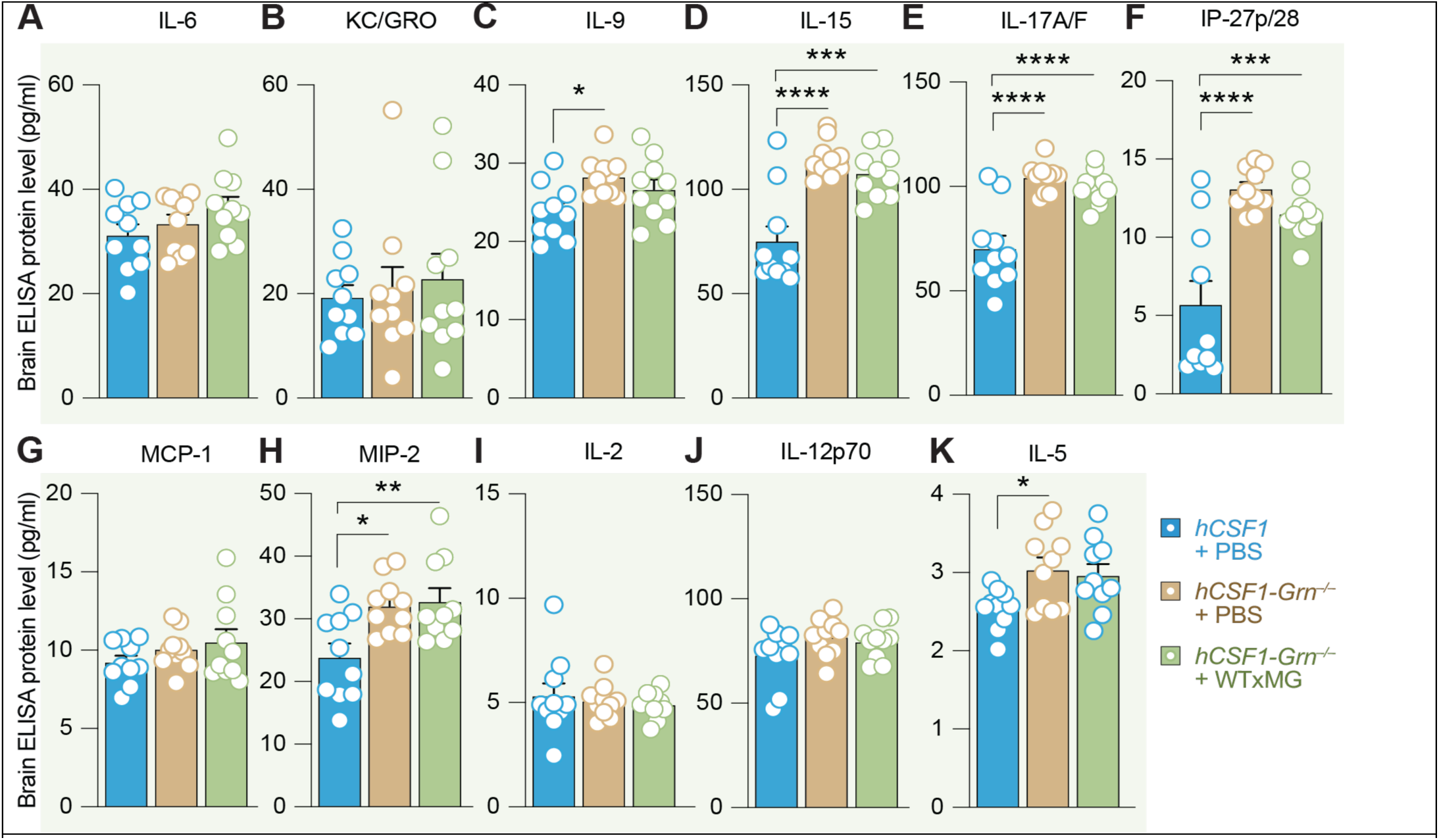
Mouse cytokine levels in the brain. A multiplex ELISA was used to concurrently examine protein levels of several murine cytokines and chemokines within the brains of *hCSF1*-PBS, *hCSF1-Grn^−/−^* +PBS, and hCSF1-Grn***^−/−^*** +xMG engrafted mice. For each analyte, N = 10 mice/group were compared by one-way ANOVA with Tukey’s multiple comparisons test. No significant (ns) differences in IL-6 **(A)** or KC/GRO **(B)** were detected. **(C)** IL-9 levels in the brain, \**p* = 0.018. **(D)** IL-15 levels (F_2,27_ = 1.238, \*\*\**p* = 0.0002 and \*\*\*\**p* < 0.0001). **(E)** IL-17A/F levels in brain (F_2,27_ = 3.026, \*\*\*\**p* < 0.0001). **(F)** IP-27p/28 levels in brain (F_2,27_ = 3.685, \*\*\**p* = 0.0006 and \*\*\*\**p* < 0.0001). **(G)** MCP-1 levels in brain, ns = nonsignificant. **(H)** MIP-2 levels in brain (F_2,27_ = 1.12, \**p* = 0.0156 and \*\**p* = 0.0083). **(I)** IL-2 levels in brain, ns = nonsignificant. **(J)** IL-12p70 levels in the brain, ns = nonsignificant. **(K)** IL-5 levels in the brain (F_2,27_ = 2.819, \**p* = 0.0367).

**Figure S3.**
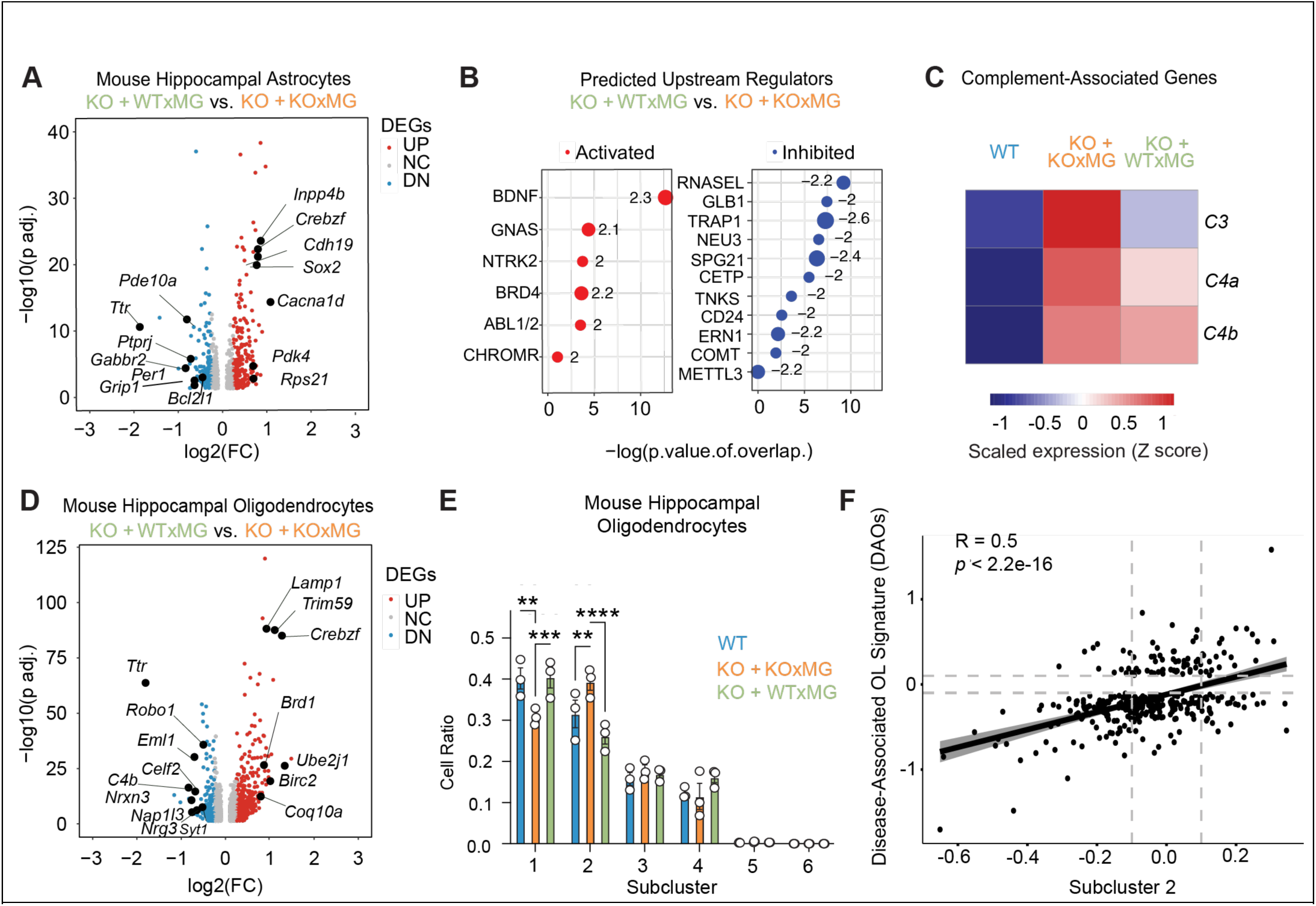
Xenografted human KOxMG or WTxMG exert differential effects on mouse hippocampal astrocyte and oligodendrocyte transcriptomes. **(A)** Volcano plot of pseudobulk differentially expressed genes (DEGs) in hippocampal astrocytes between WTxMG and KOxMG groups shows significant upregulation and downregulation of multiple genes in the KOxMG group. **(B)** Top predicted upstream regulators identified using Ingenuity Pathway Analysis (IPA). **(C)** Heatmap of complement related genes. **(D)** Volcano plot of pseudobulk DEGs in hippocampal oligodendrocytes between WTxMG and KOxMG groups highlights significant transcriptional differences in the KOxMG group. **(E)** Quantification of average cell ratios in each subcluster by group in hippocampal oligodendrocytes. **(F)** Correlation between Subcluster 2 and Disease Associated Oligodendrocyte genes, R=0.5, p<2.2 e-16.

**Figure S4.**
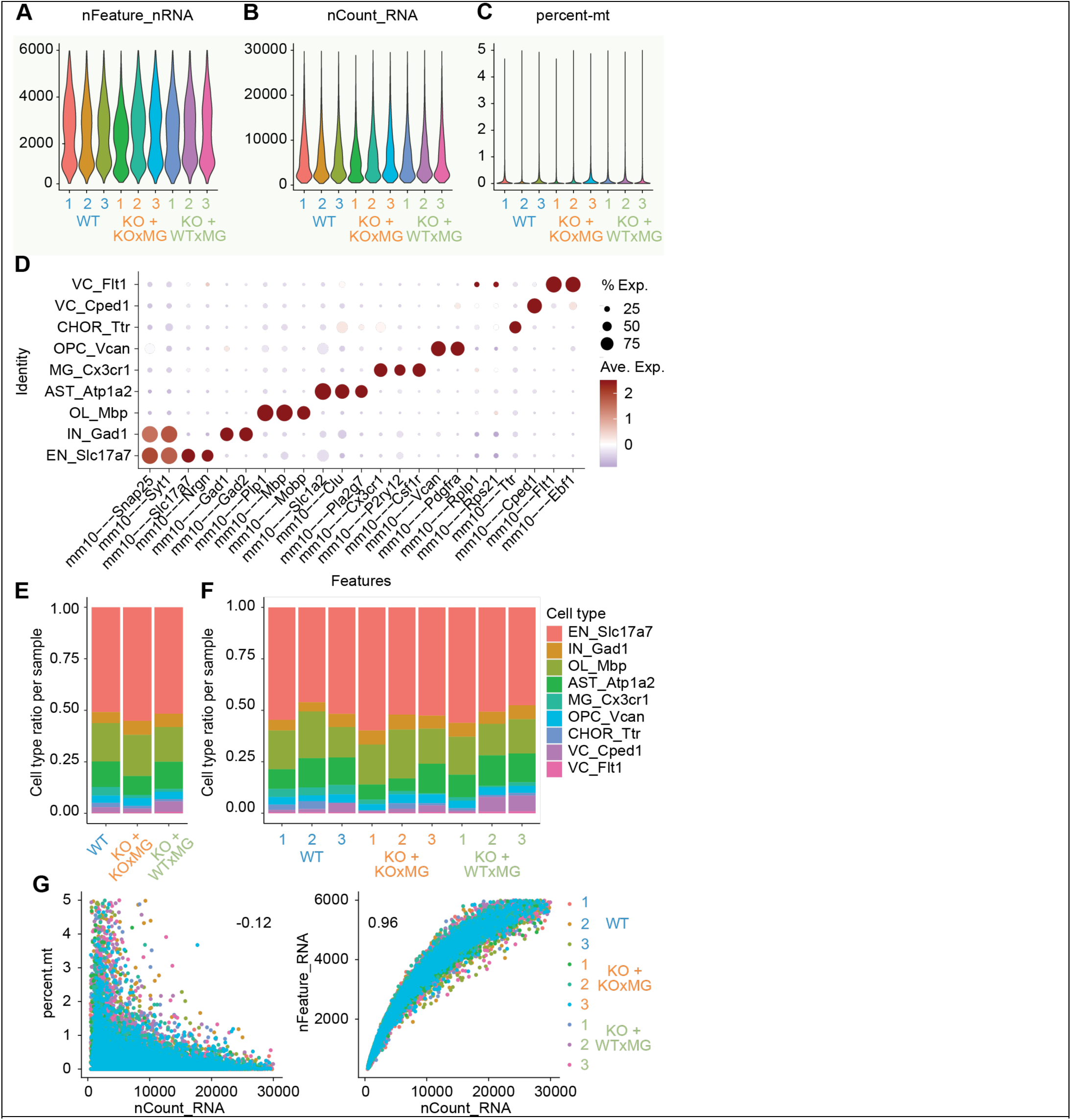
QC of snRNAseq analysis of hippocampus engrafted with *GRN^−/−^* or *GRN^+/+^* human microglia. **(A), (B)** Quality-control plots showing equivalent amounts of total RNA features and total number of RNA counts, respectively. **(C)** Violin plot showing percentage of mitochondrial genes detected per nucleus for each individual sample. **(D)** Dot plot showing expression of identity markers for each cell type. **(E)** Proportion of each cell type within each genotype. **(F)** Proportion of each cell type within individual samples. **(G)** Correlation between UMI counts and percentage of mitochondrial genes detected (left) or total gene counts (right) per nucleus for each individual sample.

**Figure S5.**
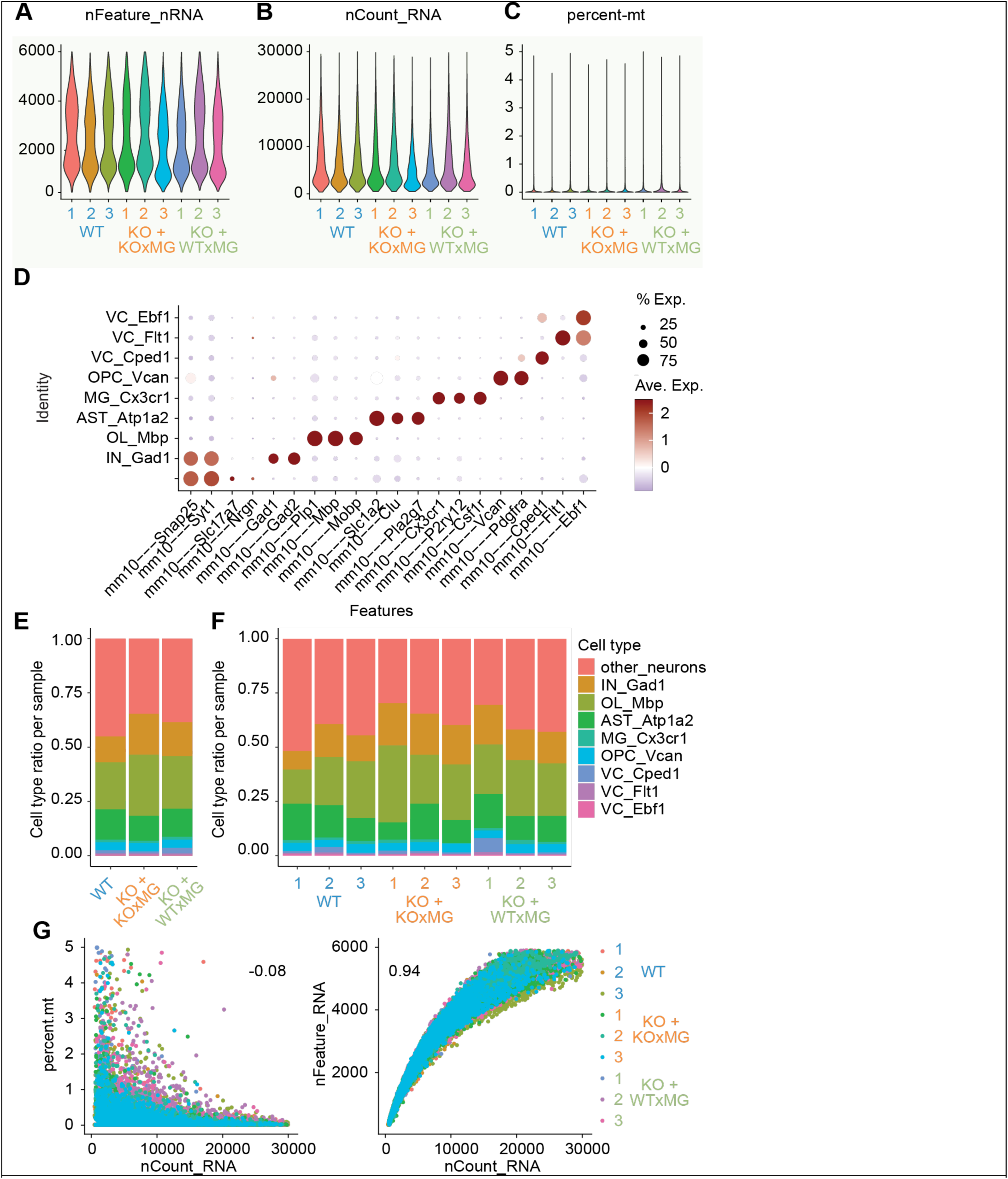
QC of snRNAseq analysis of thalamus engrafted with *GRN^−/−^* or *GRN^+/+^* human microglia. **(A), (B)** Quality-control plots showing equivalent amounts of total RNA features and total number of RNA counts, respectively. **(C)** Violin plot showing percentage of mitochondrial genes detected per nucleus for each individual sample. **(D)** Dot plot showing expression of identity markers for each cell type. **(E)** Proportion of each cell type within each genotype. **(F)** Proportion of each cell type within individual samples. **(G)** Correlation between UMI counts and percentage of mitochondrial genes detected (left) or total gene counts (right) per nuclei for each individual sample.

## Funding

This work was supported by:

DISC2-11165 (to L.G),

CIRM RT3-07893, CIRM EDUCA4-12822 (to S.K.S.)

Rainwater Charitable Foundation (to L.G.)

NIH AG048099, NIH AG055524, NIH AG056303, NIH U19 AG06970101, NIH R01NS121287 (J.T.P.), NIH AG061895 (to H.D.),

NOMIS-Gladstone fellowship and ADRC REC Scholarship (to Y.V.)

Cure Alzheimer’s Fund (to M.B-J.)

NIA T32 AG073088, T32 AG000096, Larry L. Hillblom Foundation #2025-A-175-FEL (to JP.C.)

BrightFocus Postdoctoral Fellowship Program in Alzheimer’s # A2025007F (G.E.-S.)

Alzheimer Society of Canada postdoctoral fellowship Discovery 23-10

Larry L. Hillblom Foundation #2024-A-023-FEL (to G.E.-S.).

## Authors’ contributions

Conception and design: LG, MBJ, JP, HD, SN, YV

Data acquisition: HD, YV, SN, JLG, VD, JBF, JPC, JKC, GES, AA, JN, ALC, SKS, MYW, LF, SG

Data analysis and interpretation: YV, JLG, VD, JBF, JP, HD, JPC, GES, SKS, MBJ, SN, LG, LF

Drafting of manuscript: HD, SN, YV, JP, MBJ, LG

Editing of the manuscript: YV, JLG, VD, JBF, JP, HD, JPC, JKC, GES, AA, JN, ALC, SKS, MBJ, LG

All authors approved before submission

## Competing Interests

M.B.J. is a co-founder and consultant for Savanna Biotherapeutics (formerly NovoGlia Inc.). L.G. is founder and equity holder of Aeton Therapeutics, co-founder and equity holder of NeuroVanda Therapeutics; scientific advisor of Arvinas, NeuroLamda, and consultant for Retro Biosciences.

## Data Availability

The RNA-seq data sets are available through the GEO SuperSeries accession number is currently pending. Further information and requests for resources and reagents should be directed to and will be fulfilled by Drs. Li Gan, Mathew Blurton-Jones, and Jeanne Paz.

